# Pooled CRISPR interference screening identifies crucial transcription factors in gas-fermenting *Clostridium ljungdahlii*

**DOI:** 10.1101/2024.02.20.581120

**Authors:** Huan Zhang, Huibao Feng, Xin-Hui Xing, Weihong Jiang, Chong Zhang, Yang Gu

## Abstract

Gas-fermenting *Clostridium* species hold tremendous promise for one-carbon biomanufacturing. To unlock their full potential, it is crucial to unravel and optimize the intricate regulatory networks that govern these organisms; however, this aspect is currently underexplored. In this study, we employed pooled CRISPR interference (CRISPRi) screening to uncover a wide range of functional transcription factors (TFs) in *Clostridium ljungdahlii*, a representative species of gas-fermenting *Clostridium*, with a special focus on the TFs associated with the utilization of carbon resources. Among the 425 TF candidates, we identified 75 and 68 TF genes affecting the heterotrophic and autotrophic growth of *C. ljungdahlii*, respectively. We directed our attention on two of the screened TFs, NrdR and DeoR, and revealed their pivotal roles in the regulation of deoxyribonucleotides (dNTPs) supply, carbon fixation, and product synthesis in *C. ljungdahlii*, thereby influencing the strain performance in gas fermentation. Based on this, we proceeded to optimize the expression of *deoR* in *C. ljungdahlii* by adjusting its promoter strength, leading to improved growth rate and ethanol synthesis of *C. ljungdahlii* when utilizing syngas. This study highlights the effectiveness of pooled CRISPRi screening in gas-fermenting *Clostridium* species, expanding the horizons for functional genomic research in these industrially important bacteria.

## Introduction

Microbial conversion of one-carbon (C1) gases into value-added products has provided an alternative to the traditional biomanufacturing approach that normally uses starchy feedstocks and sugars (Bae et al., 2022). Gas-fermenting *Clostridium* species is a group of chemoautotrophic bacteria that can capture CO_2_ and CO through the Wood-Ljungdahl pathway (WLP) (Liew et al., 2016), and further use these two C1 gases to produce multiple native chemicals (Abrini et al., 1994; Kopke et al., 2011). Thus, clostridial gas fermentation has shown great potential in C1 gas utilization (Liew et al., 2022; Zhang et al., 2020b). To improve the performance of these autotrophic *Clostridium* species in gas fermentation, a range of metabolic engineering strategies have been attempted in recent years (Lee et al., 2022). However, the breakthrough to achieve superior phenotype of gas-fermenting *Clostridium* strains still remains to be expected.

Metabolic networks in microorganisms are intensively regulated at different levels, in which transcriptional regulation often plays a vital role (Browning and Busby, 2004; Imam et al., 2015). Thus, to unlock the full potential and guide the improvement of industrial microorganisms, it is crucial to understand the metabolic regulation mechanism and then incorporate the related information into strain design and modification. However, to date, this aspect remains poorly explored in gas-fermenting *Clostridium* species, with only a few transcription factors that have been identified (Liu et al., 2022b; Zhang et al., 2021; Zhang, 2023; Zhang et al., 2020a). The lack of knowledge regarding crucial regulatory elements as well as the underlying regulatory mechanisms has impeded the improvement of these industrially important bacteria.

To address the above issue, we focused on functional genomic approaches (Przybyla and Gilbert, 2022). A recent study has used transposon sequencing (Tn-seq) (van Opijnen et al., 2009) to identify essential genes in gas-fermenting *Clostridium autoethanogenum* (Woods et al., 2022). This work, despite lacking detailed functional analysis, strongly supports a continued large-scale screening of crucial genes in gas-fermenting *Clostridium* strains. However, Tn-seq depends on random transposon insertions into the chromosome, usually favoring specific “hot spots” and exhibiting an insertion preference towards larger genes (Wang et al., 2018); additionally, Tn-seq is not applicable to investigate the function of a specific subset of genes of interest. As an alternative, pooled CRISPR interference (CRISPRi) screening enables genotype-phenotype association in a massively parallel manner (Bock et al., 2022), especially suitable for targeting a specific gene library of interest (Cain et al., 2020), The effectiveness of this approach in functional genomics study has been demonstrated in various model bacteria, including *Escherichia coli*, *Bacillus subtilis*, *Streptococcus pneumonia*, *Corynebacterium glutamicum*, and *Corynebacterium pyogenes* (Wang et al., 2018; Jiang et al., 2020; Liu et al., 2022a; Liu et al., 2017; Peters et al., 2016). Nevertheless, it remains unknown whether this strategy can be applied to gas-fermenting clostridia given the great differences in genetic manipulation of heterotrophic and autotrophic bacteria.

In this study, we performed pooled CRISPRi screening with a high-quality crRNA library targeting 425 TFs in gas-fermenting *Clostridium ljungdahlii*. These TFs are anticipated to regulate a multitude of genes, including those associated with C1 gas metabolism. By comparing the composition of the crRNA library before and after culturing in media containing various carbon sources as well as the subsequent experimental validation, we identified multiple previously uncharacterized TFs that could significantly affect the growth and product synthesis of *C. ljungdahlii*. Based on these data, we specially focused on two TFs, NrdR and DeoR, and revealed how they affected the *C. ljungdahlii*’s performance in gas fermentation. A further modulation of the *deoR*’s expression effectively enhanced cell growth and ethanol synthesis, highlighting the utility of this TF in driving phenotypic improvement. Our study expanded the pooled CRISPRi screening method to gas-fermenting clostridia, showcasing the effectiveness of this tool for rapid mining of the TF genes linked to important phenotypes of these gas-fermenting bacteria.

## Results

### Design and construction of a CRISPRi library targeting TFs in *C. ljungdahlii*

To discover new TFs associated with the growth of *C. ljungdahlii* on different carbon sources, we designed a library of 4,553 crRNAs for CRISPRi screening. This library includes 4,153 crRNAs targeting 425 TF genes as well as 400 negative control crRNAs with no binding hits in the *C. ljungdahlii* genome (Supplementary Data 1 and 2). For each gene, a maximum of 10 crRNAs were meticulously designed to specifically target the template strand of its open reading frame (ORF) (Zhang et al., 2017). The design process prioritized incorporating as many crRNA targeting positions as feasible within the initial 20% region of ORFs, aiming to maximize the repression activity (Methods, Fig. 1b).

**Fig. 1.**
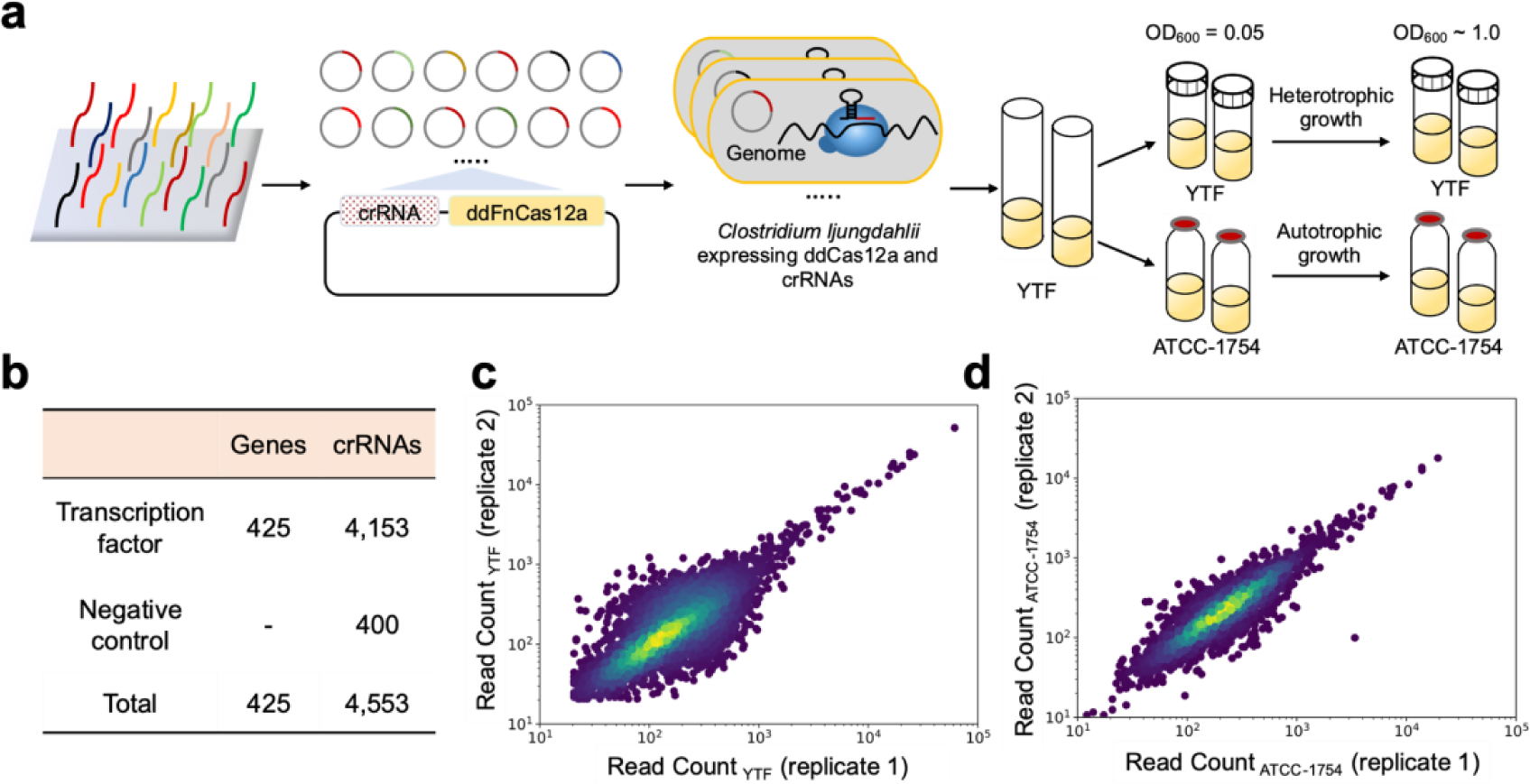
Pooled CRISPRi screening generated high-quality data for identifying TFs associated with the growth of *C. ljungdahlii* on various carbon sources. **a**, Schematic illustration of the screening protocol in *C. ljungdahlii*. A crRNA library targeting 425 TFs were synthesized on a DNA microarray. Next, oligonucleotides were amplified and integrated into expression plasmids. The plasmids were transferred into *C. ljungdahlii*, yielding cell library to be cultured under selective conditions. The abundance of each crRNA before and after cultivation was quantified via NGS. YTF, the yeast extract-tryptone-fructose medium for heterotrophic growth. ATCC-1754, the medium used for gas fermentation. **b,** The brief outline of the crRNA library. c and d, The normalized read count (see Materials and Methods) between the two biological revealed strong correlation replicates when cultivated in the YTF medium (c, heterotrophic growth, Pearson’s r = 0.7774) and the ATCC-1754 medium (d, autotrophic growth, Pearson’s r = 0.9013).

The library construction was based on our previously reported ddCas12a-based CRISPRi system for *C. ljungdahlii* (Zhao et al., 2019). To initiate the construction process, an oligo library was synthesized via DNA microarray. Subsequently, the synthesized oligonucleotides underwent PCR amplification, and the resulting DNA fragments were integrated into the expression plasmid that contained ddCas12a (Fig. 1a). Afterwards, all the plasmids were transformed into *E. coli* for amplification, and then subjected to quality evaluation via next-generation sequencing (NGS) (Fig. 1a). The result showed that the relative abundance of the majority of crRNAs (89.5%) and genes (median crRNA read count, 99.8%) was kept within 10-fold range (Supplementary Fig. 1), indicating the uniformity of the plasmid library. Such a high-quality plasmid library was then extracted from *E. coli* and transformed into *C. ljungdahlii* by electroporation, yielding a cell library containing ∼250,000 transformants with an approximately 50-fold coverage. Subsequently, the cell library was cultured in the YTF (yeast extract-tryptone-fructose) medium and the modified ATCC-1754 medium (using CO_2_/CO as the carbon sources) to screen for TF genes associated with the heterotrophic and autotrophic growth of *C. ljungdahlii*, respectively. When OD_600_ of the grown cells reached ∼1.0, they were collected. The crRNA coding regions of each cell were PCR amplified and quantified through NGS (Fig. 1a). As expected, a high consistency was observed between the two biological replicates, indicating the reliability of the screening experiments (Figs. 1c and 1d).

### CRISPRi screening of TFs associated with heterotrophic growth of *C. ljungdahlii*

We subsequently employed the CRISPRi library for rapid screening of TFs associated with the heterotrophic growth of *C. ljungdahlii* (Supplementary Fig. 2). Briefly, the transformed *C. ljungdahlii* cells were transferred into the nutrient-rich YTF medium with an initial OD_600_ of ∼ 0.05 and cultivated until reaching an OD_600_ of ∼ 1.0. Then, the grown cells were subcultured in the same medium again. Two independent biological replicates were used to test the variability between samples (Fig. 1c). Next, both the seed culture and the subculture were collected for NGS, aiming to compare their difference in crRNA abundance. Following data analysis, we discovered that 75 TFs had potential association with the heterotrophic growth of *C. ljungdahlii*, in which ten candidates resulted in enhanced cell growth after CRISPRi-based repression, while the others impaired cell growth (Score ≥ 0.65) (Fig. 2a and Supplementary Data 3).

**Fig. 2.**
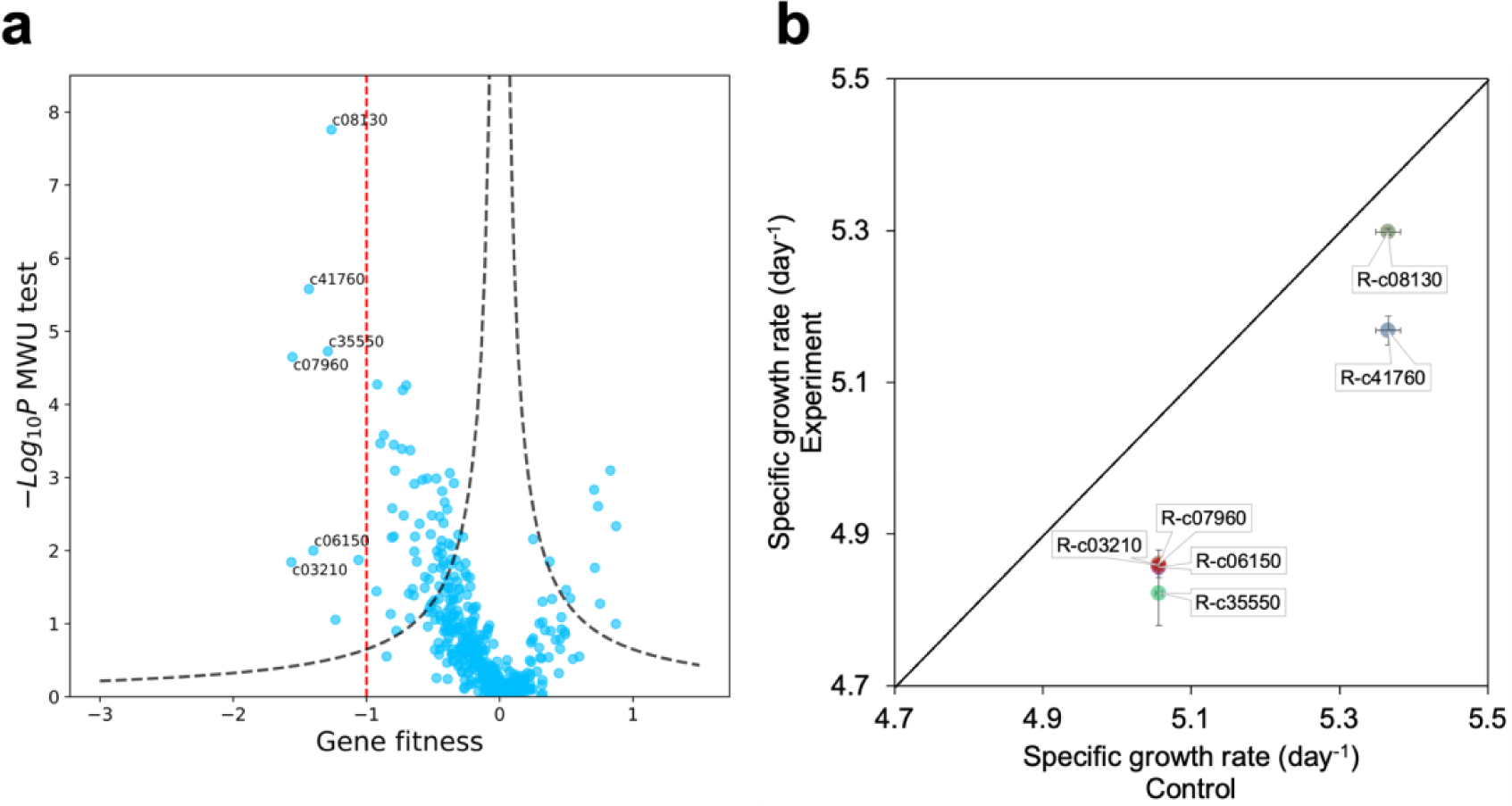
Identification of the TFs linked to the heterotrophic growth of *C. ljungdahlii*. **a**, A volcano plot of gene fitness versus −log_10_*P* (two-tailed MWU test). The threshold for discerning functional genes (Score = 0.65) was indicated by the black dashed line. Six TF genes with fitness below −1 (indicated by the red dashed line) were selected for individual validation. **b**, The growth changes of *C. ljungdahlii* caused by the repression of the abovementioned six TF genes with fitness below −1. Compared with the control strain (lacking crRNA), the repression of these six TF genes showed significantly lower growth rate during the logarithmic phase (the initial 24 h of cultivation). The error bars indicated the standard deviation across three biological replicates (*n* = 3).

To validate the above screening result, we selected six TF genes (CLJU_c41760, CLJU_c08130, CLJU_c07960, CLJU_c06150, CLJU_c35550, and CLJU_c03210) with fitness values below −1 for individual knockdown using CRISPRi and then examined phenotypic outcomes. As expected, the repression of these genes did change the growth of *C. ljungdahlii* compared to the control strain (containing a plasmid carrying ddCas12a but without crRNA), in which CLJU_c41760 and CLJU_c08130 significantly impaired the growth rate and final biomass, while the other four genes only caused slightly slower growth rate but no substantial disparity of the final biomass (Fig. 2b and Supplementary Fig. 3). Together, Therefore, these findings demonstrated the validity of our screening experiment.

### CRISPRi screening of TFs associated with the growth of *C. ljungdahlii* on syngas

The real potential of *C. ljungdahlii* lies in its capacity to efficiently utilize C1 gases such as CO and CO_2_ for bioproduction. In light of this, we attempted to employ the CRISPRi library for quickly identifying TFs capable of affecting C1 gas metabolism in *C. ljungdahlii*. As shown in Supplementary Fig. 4, the seed culture was prepared using a consistent approach, wherein 2 mL of the *C. ljungdahlii* transformants were incubated in 600 mL of the YTF medium. When cell density reached OD_600_ of ∼ 1.0, the grown cells (1.5 ml) were transferred into 30 mL of the ATCC-1754 medium for gas fermentation (syngas: CO_2_/CO/H_2_/N_2_) with two biological replicates (Fig. 1d). After the OD_600_ of the culture reached ∼ 1.0 again, cells were collected for NGS.

Based on the NGS data, we assessed the variation in abundance of each crRNA before and after the cultivation in syngas, which was further compared with the results of the negative control. Following a rigorous statistical analysis, we identified a total of 68 TFs (Score ≥ 0.65) that are potentially associated with the growth of *C. ljungdahlii* on syngas. Among these TF genes, 66 led to the impaired cell growth after transcriptional repression, while the remaining two promoted cell growth after repression (Fig. 3a and Supplementary Data 4). Interestingly, 33 out of the 68 TF genes also occurred in those found to influence the heterotrophic growth of *C. ljungdahlii* (Fig. 3c and Supplementary Data 5), implying that these 33 TF genes play crucial regulatory roles in *C. ljungdahlii* regardless of carbon sources. In contrast, the remaining 35 TF genes appear to be specifically associated with the growth of *C. ljungdahlii* on C1 gases (Supplementary Data 6).

**Fig. 3.**
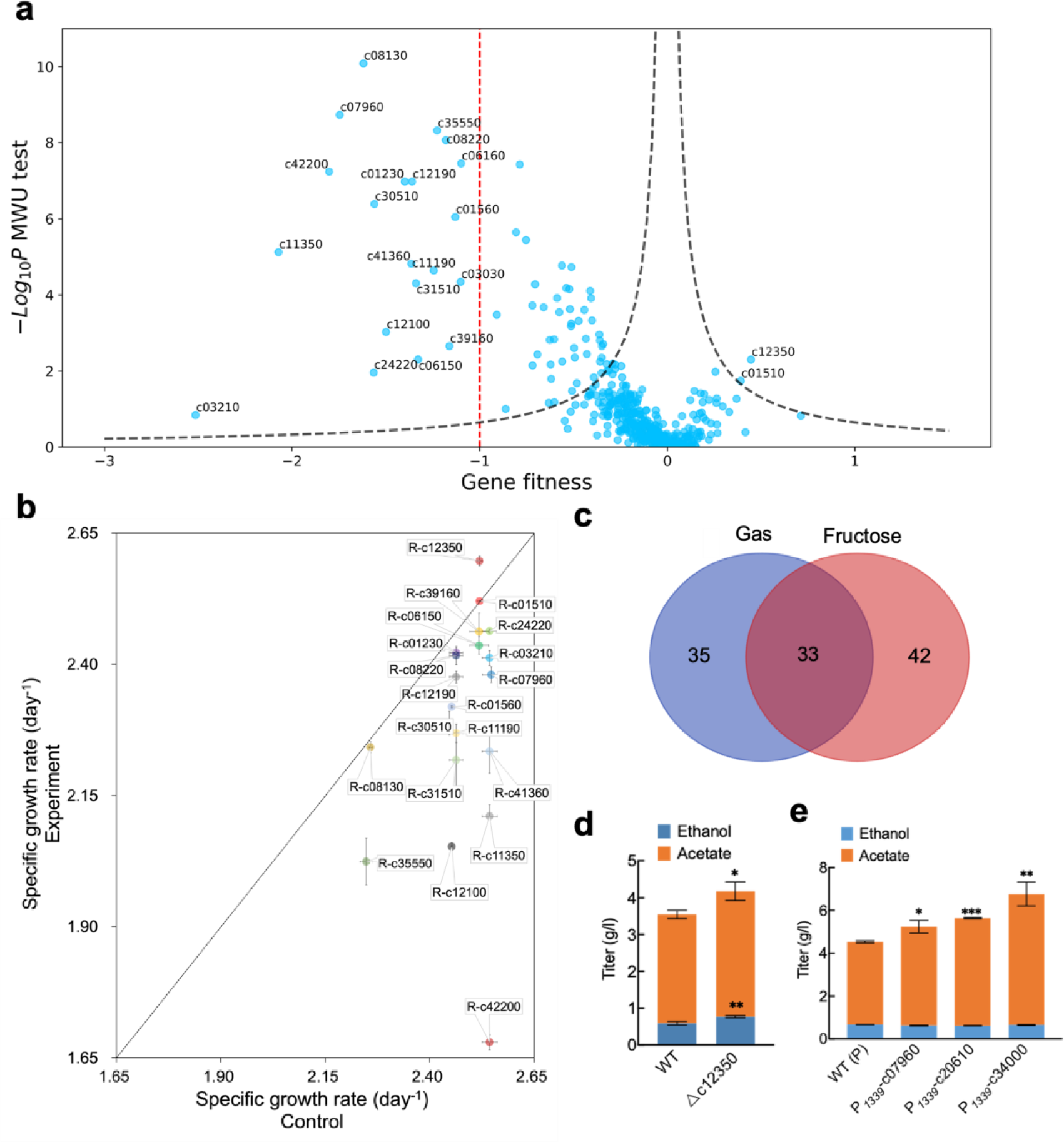
Identification of the TFs linked to the autotrophic growth of *C. ljungdahlii* on syngas. **a**, A volcano plot of gene fitness score (Gene fitness) versus −log_10_*P* (two-tailed MWU test). The threshold for discerning functional genes (Score = 0.65) was indicated by the black dashed line. 18 TFs with fitness below −1 (indicated by the red dashed line) and 2 TFs with fitness above 0.35 were selected for individual validation. **b,** The growth changes of *C. ljungdahlii* caused by the repression of the abovementioned 20 TF genes. The repression of CLJU_c12350 led to enhanced growth rate, while the repression of CLJU_c01510 and CLJU_c08130 had no influence on cell growth. In contrast, the repression of the remaining 17 TF genes decreased growth rate. The growth rate was calculated based on the growth profile in the logarithmic phase (the initial 48 h of cultivation). The error bars indicated the standard deviation across three biological replicates (*n* = 3). **c,** The Venn diagram that showcases the overlap between two screening outcomes based on different carbon sources (fructose and syngas). **d,** The influence of the deletion of CLJU_c12350 (*nrdR*) on the production of acetate and ethanol by *C. ljungdahlii* in gas fermentation. WT, the wild-type strain. Statistical analysis was performed based on the two-tailed Student’s *t*-test. Significant levels were denoted as: *, *P* < 0.001; **, *P* < 0.01; versus WT. **e**, The influence of overexpressing CLJU_c20610 (*deoR*), CLJU_c07960, and CLJU_c34000 on acetate formation. Data are presented as mean ± standard deviation (*n* = 3). Statistical analysis was performed based on the two-tailed Student’s *t*-test. Significant levels were denoted as: *, *P* < 0.05; **, *P* < 0.01; ***, *P* < 0.001; versus WT.

To validate the reliability of the above screening results, we chose 20 TF genes that generated substantial changes in cell growth (Score ≥ 0.65, 2 with Fitness ≥ 0.35 and 18 with Fitness ≤ −1) for a further investigation. After individual repression, most of these TF genes (18/20) gave the similar changes in cell growth rate as observed in the aforementioned screening of CRISPRi library (Fig. 3b and Supplementary Fig. 5), with the exception of the CLJU_c01510 and CLJU_c08130 genes, which did not significantly change the growth rate of *C. ljungdahlii* upon repression (Fig. 3b and Supplementary Fig. 5). These 18 TF genes are distributed in the HrcA, RpoD, ArgR, Fur2, and CtsR families (Supplementary Data 7). Hence, we confirmed the reliability of the screening results regarding the TFs involved in regulating the growth of *C. ljungdahlii* on C1 gases.

Next, we proceeded to investigate if these TF genes can be used as genetic modification targets to improve the performance of *C. ljungahlii* in gas fermentation. Specifically, the following genetic manipulations were conducted to examine phenotypic outcomes: (i) 16 TF genes (CLJU_c07960, CLJU_c20610, CLJU_c34000, <colcnt=5> CLJU_c11350, CLJU_c42200, CLJU_c41360, CLJU_c31510, CLJU_c14130, CLJU_c34650, CLJU_c12100, CLJU_c35550, CLJU_c30510, CLJU_c28220, CLJU_c32060, CLJU_c01860, and CLJU_c37800) were chosen for overexpression because their potential negative impact on the growth of *C. ljungdahlii* after repression (Score ≥ 0.65), as indicated by the screening data (Fig. 3a). (ii) the CLJU_c12350 gene was deleted because its repression resulted in the improved growth of *C. ljungdahlii* (Fig. 3b). The yielding 17 strains were cultivated in the ATCC-1754 medium for gas fermentation, yielding four engineered strains that exhibited improved ability in product synthesis compared with the control strain (Figs. 3d and 3e). Specifically, the deletion of CLJU_c12350 led to increased production of both acetate and ethanol (Fig. 3d), while overexpressing CLJU_c20610, CLJU_c07960, or CLJU_c34000 resulted in elevated acetate formation exclusively (Fig. 3e). In contrast, the manipulation of the remaining 13 TF genes did not cause significant alteration in product formation (Supplementary Fig. 6).

### Elucidating the regulatory role of NrdR in *C. ljungdahlii*

The abovementioned CLJU_c12350 gene is annotated to encode the regulator NrdR in *C. ljungdahlii*. NrdR has been known to negatively regulate the expression of ribonucleotide reductase (RNR) genes (including *nrdAB*, *nrdJ*, and *nrdD*) in some heterotrophic bacteria (Grinberg et al., 2022). The *nrd* genes are responsible for the production of dNTPs, and thus, the metabolic regulation of these genes is crucial for maintaining a balanced pool of dNTPs for DNA replication and repair in cells (Crespo et al., 2015). To investigate whether the *C. ljungdahlii* NrdR (*Clju*NrdR) also regulates the expression of *nrd* genes, we deleted CLJU_c12350 and then examined the transcriptional variation of *nrdA* (CLJU_c06840) and *nrdD* (CLJU_c01840) (Fig. 4a). As shown in Fig. 4b, the deletion of CLJU_c12350 resulted in 7.3 and 6.5-fold upregulation of *nrdA* and *nrdD*, respectively, indicating that *Clju*NrdR can regulate these two *nrd* genes in *C. ljungdahlii*.

**Fig. 4.**
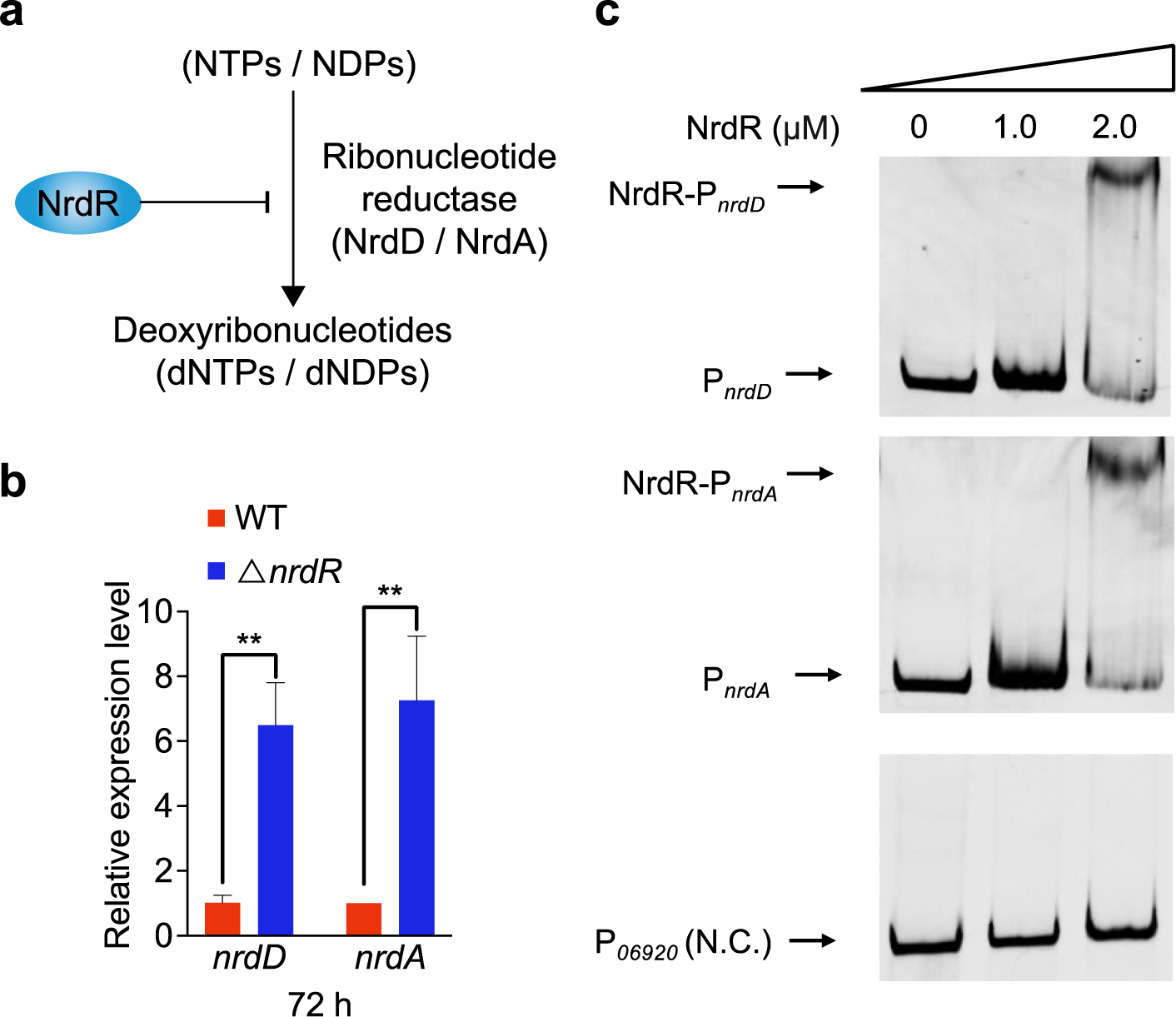
The *Clju*NrdR’s regulation on dNTPs biosynthesis in *C. ljungdahlii*. **a**, The control of *Clju*NrdR on the formation of dNTPs. **b,** The influence of the deletion of *nrdR* (encoding *Clju*NrdR) on the expression of *nrdD* and *nrdA*. Data are presented as mean ± standard deviation (*n* = 3). Statistical analysis was performed based on the two-tailed Student’s *t*-test. **, *P* < 0.01. **c,** EMSAs for identifying the interaction between *Clju*NrdR and *nrdD* or *nrdA*. The DNA probe containing the promoter region of the CLJU_c06920 gene was used as the negative control.

The following question is whether *Clju*NrdR can directly control the expression of *nrdA* and *nrdD*. Thus, we purified the *Clju*NrdR protein and determined its binding activities to the promoter regions of these two genes through electrophoretic mobility shift assays (EMSAs). As depicted in Fig. 4c, the presence of *Clju*NrdR at the concentration of 4 μM could generate a clear mobility shift for both DNA probes, strongly suggesting that *Clju*NrdR can directly bind to the promoter regions of *nrdA* and *nrdD* and then regulated their expression. Given that *nrdA* and *nrdD* are crucial for the synthesis of dNTPs/dNDPs (Chen et al., 2019; Grinberg et al., 2022), the direct control of *Clju*NrdR on these genes may contribute to balancing the dNTP level in *C. ljungdahlii*, consequently affecting the physiology and metabolism of this bacterium.

### Elucidating the regulatory role of DeoR in *C. ljungdahlii*

Among the aforementioned three TF genes (CLJU_07960, CLJU_20610, and CLJU_34000) that, upon overexpression, improved the product synthesis of *C. ljungdahlii* in gas fermentation (Fig. 3e), CLJU_c20610 encodes a TF belonging to the DeoR family. The DeoR family regulators have been found to regulate substrate metabolism in bacteria such as *Haloferax volcanii* (Martin et al., 2018). Thus, we speculated that the *C. ljungdahlii* DeoR (*Clju*DeoR) (encoded by CLJU_c20610) may control CO_2_/CO fixation and assimilation in gas fermentation. To confirm this possibility, we examined transcriptional changes of all the genes located in WLP, the pathway responsible for carbon fixation and assimilation in *C. ljungdahlii* (Fig. 5a), following the overexpression of CLJU_c20610. In addition, given that the CLJU_c20610 overexpression could change the product formation in *C. ljungdahlii* (Fig. 3e), the transcriptional changes of the crucial genes responsible for acetate and ethanol synthesis were also tested. As expected, three WLP genes (CLJU_c20040, CLJU_c17910, and CLJU_c37670) and one gene (CLJU_c16510) that is key for ethanol formation were found to be significantly upregulated with the overexpression of CLJU_c20610 (Fig. 5a), thereby indicating that they are under the regulation of *Clju*DeoR. To further explore whether *Clju*DeoR directly controls these genes, the protein was purified and subjected to EMSAs. As shown in Fig. 5b, *Clju*DeoR gave a clear mobility shift for all the four DNA probes at the protein concentration of 2.0 μM, demonstrating its binding activity to the promoter regions of these genes. Therefore, these results suggest the direct regulation of *Clju*DeoR on carbon fixation and assimilation as well as product synthesis in *C. ljungdahlii*, indicating its importance in this bacterium. To our knowledge, such a regulatory role of DeoR in autotrophic bacteria has not been reported previously.

**Fig. 5.**
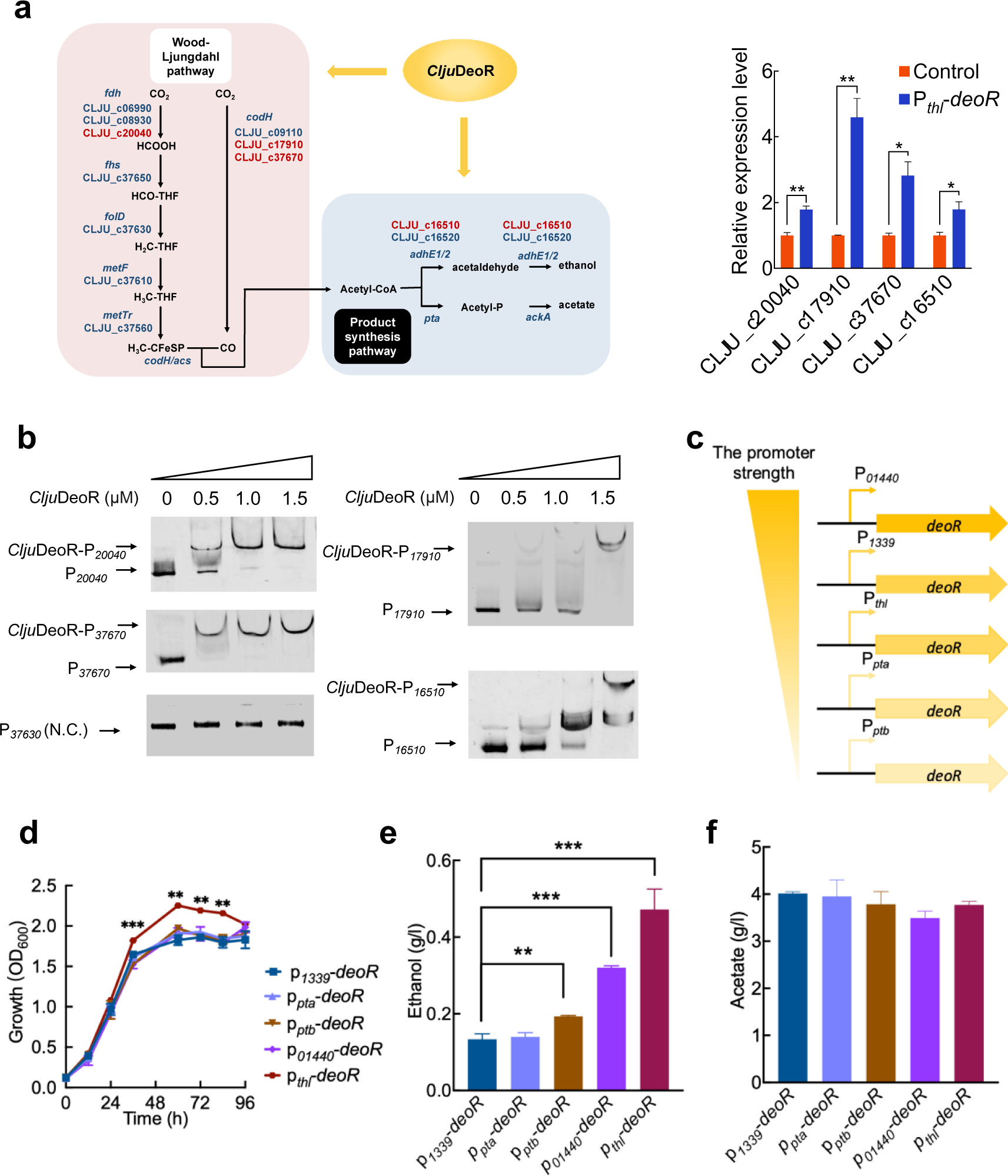
Dissection of the *Clju*DeoR’s regulatory role in *C. ljungdahlii*. **a**, The regulation of *Clju*DeoR on WLP (pink) and product synthesis pathway (cyan) in *C. ljungdahlii*. The genes that may be directly regulated by *Clju*DeoR were shown in red. *fdh*, encoding formate dehydrogenase; *fhs* encoding formyl-THF synthetase; *fold* encoding formyl-THF cyclohydrolase/methylene-THF dehydrogenase; *metF*, encoding methylene-THF reductase; *metTr*, encoding methyltransferase; *codH/acs*, encoding carbon monoxide dehydrogenase/acetyl-CoA synthase; *codH*, encoding carbon monoxide dehydrogenase; *adhE1/2*, encoding aldehyde/alcohol dehydrogenase 1 & 2; *pta*, encoding phosphotransacetylase; *ackA*, encoding acetate kinase. Genes that exhibited significant upregulation following the deletion of *deoR* were highlighted in red. Data are presented as mean ± standard deviation (*n* = 3). Statistical analysis was performed using the two-tailed Student’s *t*-test. *, *P* < 0.05; **, *P* < 0.01. **b,** EMSA for evaluating the binding affinity of *Clju*DeoR to the promoter region of the CLJU_c20040, CLJU_c17910, CLJU_c37670, and CLJU_c16510 genes. The DNA probe containing the promoter region of the CLJU_c37360 gene was used as the negative control. **c,** Fine-tuning the expression of *deoR* using different promoters. The strength of the promoters was denoted by color depth. **d,** The growth curves of the *C. ljungdahlii* strains overexpressing the *deoR* gene with the P*_01440_*, P*_pta_*, P*_ptb_*, and P*_thl_* promoters in gas fermentation. The strain overexpressing *deoR* with the P*_1339_* promoter was used as the control. Data are presented as mean ± standard deviation (*n* = 3). Statistical analysis was performed using the two-tailed Student’s *t*-test. **, *P* < 0.01; ***, *P* < 0.001; versus P*_1339_*-*deoR*. **e and f,** The production of ethanol (**e**) and acetate (**f**) by the abovementioned *deoR*-overexpressing *C. ljungdahlii* strains in gas fermentation. Data are presented as mean ± standard deviation (*n* = 3). Statistical analysis was performed using the two-tailed Student’s *t*-test. **, *P* < 0.01; ***, *P* < 0.001; versus P*_1339_*-*deoR*.

Given the importance of *Clju*DeoR as well as its promotion on the product synthesis through gene overexpression in *C. ljungdahlii* (Fig. 3e), we attempted to optimize the positive regulatory effect of *Clju*DeoR by adjusting its expressional level in cells. To this end, multiple promoters, which exhibited varying activities in *C. ljungdahlii* (Huang et al., 2016; Zhao et al., 2019), were utilized to drive the expression of CLJU_c20610 (Fig. 5c). Among these promoters, P*_thl_* generated a significantly enhanced cell growth compared to the control (the original promoter P*_1339_*), while the other promoters had no discernible promotion on cell growth (Fig. 5d). Meanwhile, an increased ethanol production was observed with the use of the P*_thl_*, P*_01440_*, and P*_ptb_* promoters, in which P*_thl_* gave the highest increase (3.17-fold) (Fig. 5e); in contrast, no substantial change was observed in acetate production by replacing the promoter (Fig. 5f). Interestingly, P*_thl_* that led to the highest increase in cell growth and ethanol formation of *C. ljungdahlii* was not the strongest one among the tested promoters (Fig. 5c). Therefore, these findings suggest that an appropriate expression level of *Clju*DeoR is crucial for optimizing its regulation in *C. ljungdahlii*.

## Discussion

In this study, we employed pooled CRISPRi screening, a high-throughput screening method not yet applied to autotrophic *Clostridium* species, to identify crucial TF genes associated with the growth and product synthesis in gas-fermenting *C. ljungdahlii*. With a particular focus on two crucial TFs, *Clju*NrdR and *Clju*DeoR, we conducted additional analyses to gain insights into their regulatory roles in *C. ljungdahlii*. These results demonstrate the effectiveness of pooled CRISPRi screening in uncovering functional genes relevant to important phenotypes of autotrophic bacteria.

Our list of screened TFs provides an important resource store for further studies into the regulatory mechanisms associated with the performance of *C. ljungdahlii*. Actually, in addition to *Clju*NrdR and *Clju*DeoR, some other screened TFs also showed potential importance in *C. ljungdahlii* (Fig. 3a). For example, the overexpression of CLJU_c07960 could significantly enhance the production of acetate (Fig. 3e), although acetate is usually not the target product of *C. ljungdahlii*. The CLJU_c07960 gene is predicted to encode a heat shock regulatory protein HrcA, which has been found to control the expression of class I stress genes including *groES*, *groEL*, *grpE*, *dnaK*, *dnaJ*, and *htpG* in bacteria (Schulz and Schumann, 1996; Selby et al., 2011; Suo et al., 2017). The presence of HrcA can maintain the expression of these stress genes at a relatively low level; but such a repression can be released with the exposure to stress (Schumann, 2003). However, the function of HrcA in *Clostridium acetobutylicum*, an important solventogenic *Clostridium* species (Wang et al., 2013), is likely different from the findings in some well-studied microorganisms such as *Bacillus subtilis* and *Staphylococcus aureus* (Chastanet et al., 2003; Schumann, 2003). These studies indicate the complexity of the HrcA’s function in microorganisms, and thus, its regulatory role as well as the underlying mechanism in *C. ljungdahlii* merits further explorations. It is worth noting that the phenotypic consequences of CRISPRi are highly relied on the repression dosage (Hawkins et al., 2020). Despite our efforts in designing 10 crRNAs for each gene and employing statistical analyses to mitigate this issue, it remains possible that some functional genes were undiscovered in the screening experiments due to the absence of highly active crRNAs. Therefore, a more comprehensive validation may be necessary to investigate genes that were not identified as functional in this study.

In addition, we also noticed that, among the TFs subjected to phenotypic analysis, the ratio of the TFs that could generate improved product synthesis was low (Fig. 3a). This may be attributed to: (i) according to the NGS data (Fig. 3a), the vast majority of the TFs decreased the growth of *C. ljungdahlii* after their repression by CRISPRi. This finding indicates the importance of these TFs for *C. ljungdahlii*’s growth in gas fermentation but does not mean that they must be positively correlated with cell growth or product synthesis. (ii) typically, TFs are capable of controlling the expression of multiple target genes. Therefore, their overexpression will generate pleiotropic or global effects, causing either improved or impaired cellular performance, which may ultimately offset each other. Therefore, fine-tuning the expression of these TFs may generate more positive phenotypic changes.

A further direction for advancing this research is to expand the size of the screening library to the whole genome scale, which would enable an investigation of all the genes associated with important phenotypes of *C. ljungdahlii*. To construct a crRNA library targeting all the genes (4,184) in *C. ljungdahlii*, an estimated minimum of 8 × 10^5^ transformants would be required. This estimate is based on the assumption that 10 crRNAs are designed for each gene as well as that the number of the required transformants should be 20 times the size of the plasmid library. This goal, however, is currently being impeded by the relatively low electroporation efficiency (∼10^4^ transformants/μg plasmids) of *C. ljungdahlii*. Hence, for large-scale genomic functional study on this bacterium, a further increased transformation efficiency of exogenous plasmid DNA is necessary. Additionally, the prominent advantage of the pooled CRISPRi screening is that it can be designed to target a specific range of genes or intergenic regions of interest. Thus, a combination of CRISPRi screening and other high-throughput methods, such as Tn-seq, which targets genome randomly and has recently been developed in gas-fermenting *Clostridium* species (Woods et al., 2022), may enable a more comprehensive and thorough functional genomics study of these autotrophic bacteria.

In summary, we successfully constructed a CRISPRi library targeting all the TF genes in *C. ljungdahlii* and performed pooled screening experiments, which allowing for rapid identification of crucial TFs in this autotrophic bacterium. We anticipate that our work will pave the way for exploring genotype-phenotype associations in gas-fermenting *Clostridium* species based on either genome-wide or specific sgRNA libraries, thereby expediting the progress of functional genomics research on these industrially significant bacteria.

## Materials and Methods

### Strains and growth media

Molecular cloning was performed using *E. coli* DH5α as the host. The *E. coil* DH5α and its derived strains were cultured in LB (lysogeny broth) medium (Bertani, 1951). The *C. ljungdahlii* DSM 13528 and its derived strains were cultured in the YTF (yeast extract-tryptone-fructose) medium for inoculum preparation (Humphreys et al., 2015), and a modified ATCC-1754 medium for gas fermentation (CO-CO_2_-H_2_-N_2_: 56%-20%-9%-15%; pressurized to 0.2 MPa) (Huang et al., 2016). All the manipulations of the *C. ljungdahlii* strains were performed in an 37℃ anaerobic chamber (Whitley A35 workstation, don Whitley Scientific). Chloramphenicol (12.5 μg/ml) and thiamphenicol (5 μg/ml) were added into media when needed. All the strains used in this study are listed in Table 1.

### DNA manipulation and reagents

All the primers were synthesized by Biosune (Biosune, Shanghai, China). The crRNA library was synthesized by GENEWIZ (GENEWIZ, Suzhou, China). The restriction enzymes were purchased from Thermo Fisher Scientific (Thermo Fisher Scientific, Waltham, MA, USA). For plasmid construction, PCR amplification was performed using KOD-Plus and KOD FX DNA polymerases from Toyobo (Toyobo, Osaka, Japan). The DNA fragments were assembled using ClonExpress II One Step Cloning Kit purchased from Vazyme (Vazyme Biotech Co., Ltd., Nanjing, China). Plasmid extraction and DNA purification were performed using the AxyPrep™ Plasmid Miniprep Kit and AxyPrep™ DNA Gel Extraction Kit from Axygen (Axygen, Hangzhou, China). For NGS library preparation, PCRs were carried out using KAPA HiFi HotStart DNA polymerase from KAPA Biosystems (KAPA Biosystems, Boston, USA). The resulting PCR products were purified using the DNA Gel Extraction Kit from New England Biolab (New England Biolab, Ipswich, MA, USA).

### Plasmid construction

The plasmids and primers used in this study were listed in Supplementary Tables 1 and 2, respectively.

The pMTLCas-*nrdR* plasmid that was designed for deleting the *nrdR* gene was constructed as follows. In brief, a crRNA fragment (AATAATTTCCCCTATCTCCG) targeting the *nrdR* gene was obtained through PCR amplification using the pMTLCas-*pta* plasmid (Huang et al., 2016) as the template and the primers *nrdR*-sgRNA-for/sgRNA-rev. Then, two homologous arms (HAs) flanking the coding region of *nrdR* were generated by PCR amplification using the *C. ljungdahlii* genomic DNA as the template and the primers *nrdR*-UpArm-for/*nrdR*-UpArm-rev (for upstream HA) and *nrdR*-DownArm-for/*nrdR*-DownArm-rev (for downstream HA). Next, the sgRNA and two HA fragments were combined through overlapping PCR using the primers *nrdR*-sgRNA-for/*nrdR*-DownArm-rev, resulting in a merged DNA fragment (sgRNA-HAs). The pMTLCas-*pta* plasmid was digested with both *Sal*I and *Xho*I to obtain a linear pMTLCas vector. Finally, the sgRNA-HAs fragment was linked to the linear pMTLCas vector using the ClonExpress II One Step Cloning Kit, generating the pMTLCas-*nrdR* plasmid. The construction of the other plasmids for gene deletion in *C. ljungdahlii* were performed via the same steps with the exception of using different sgRNAs and HAs.

The pMTL83151-P*_1339_*-*deoR* plasmid for overexpressing *deoR* was constructed as follows. In brief, the pMTL83151 plasmid (Heap et al., 2009) was linearized by treatment with both *Nde*I and *Hind*III. The P*_1339_* promoter was obtained by PCR amplification using the *C. ljungdahlii* genomic DNA as the template and the primers P*_1339_*-for/P*_1339_*-rev. The DNA fragment of *deoR* was obtained by PCR amplification using the *C. ljungdahlii* genomic DNA as template and the primers *deoR* (P*_1339_*)-for/*deoR*-rev. Then, the P*_1339_* promoter and the *deoR* gene were assembled via overlapping PCR using the primers P*_1339_*-for/*deoR*-rev, generating a merged DNA fragment P*_1339_*-*deoR*. Next, the linear pMTL83151 vector and the P*_1339_*-*deoR* fragment were assembled using the ClonExpress II One Step Cloning Kit, yielding the pMTL83151-P*_1339_*-*deoR* plasmid. Similar steps were followed to construct the other plasmids for gene overexpression in *C. ljungdahlii*.

The pZG-ddFnCas12a-*nrdR* plasmid for the transcriptional repression of *nrdR* by CRISPRi was constructed as follows. In brief, a double-strand DNA (harboring the crRNA spacer and repeat sequences) (designed on https://benchling.com) was first obtained by PCR amplification using the primers *nrdR*-crRNA-for/*nrdR*-crRNA-rev. Next, this DNA fragment was used as the template for PCR amplification again with the primers Universal-for/ Universal-rev, yielding a new DNA fragment containing the crRNA sequence and a terminator. The DNA fragment was assembled with the linear pZG-ddFnCas12a plasmid (*BamH*I/*Nco*I digestion) using the ClonExpress II One Step Cloning Kit, yielding the pZG-ddFnCas12a-*nrdR* plasmid. The other plasmids for gene repression in *C. ljungdahlii* were constructed following the same steps except using different crRNAs.

The pET28a-*nrdR*/*deoR* plasmid for producing the NrdR and DeoR protein in *E. coil* BL21(DE3) was constructed as follows. In brief, the *nrdR* and *deoR* gene were obtained by PCR amplification using the *C. ljungdahlii* genomic DNA as the template and primers pET28a-*nrdR*-F/pET28a-*nrdR*-R (for *nrdR*) or pET28a-*deoR*-F/pET28a-*deoR*-R (for *deoR*). Next, the pET28a plasmid was subjected to the digestion with *Nde*I and *Xho*I. The linear pET28a vector was then assembled with *nrdR* and *deoR* using the ClonExpress II One Step Cloning Kit, yielding the pET28a-*nrdR* and pET28a-*deoR* plasmid, respectively.

### Design of the crRNA library

The *C. ljungdahlii* genome sequence and relevant gene annotations (NC_014328.1) were used for crRNA (24-mer) design. For each TF, we searched all potential targets (TTVN24) located on the non-template strand. The SeqMap package (Jiang and Wong, 2008) was applied to check potential off-target sites by searching for “TTV(CTV/YTT)N24” 27-mers in NC_014328.1 with a tolerance setting of five mismatches. Customized scoring metrics inferred from previous reports and illustrated in Supplementary Fig. 7 were designed to evaluate the potential off-target sites identified by SeqMap. Specifically, the protospacer region of potential off-target sites is divided into three parts (6, 12, and 6 nt, from the 5’ end to the 3’ end as Region I, II and III, respectively) according to the distance to the PAM site. The partitioning of regions in this method follows the previous experimental findings of Cas12a (Kim et al., 2017), namely that the initial 6 nt constitute the seed region, the middle 12 nt form the trunk region, and the last 6 nt make up the promiscuous region. If the PAM sequence of off-target hit is “TTV”, the mismatch penalties of the three regions were set as 8, 4, 1, respectively; the penalties for non-canonical PAM “CTV” and “YTT” were set as 10, 7, 3, respectively. The off-target site was considered significant when Σ(penalty × mismatch) < 20, the corresponding crRNAs were eliminated in the subsequent processing. It’s important to highlight that the scoring metrics were designed to detect sequences with a lower likelihood of off-target effects, giving priority to those with fewer mismatches in the PAM-proximal region in comparison to the target. We also restricted the GC-content of each crRNA between 20% and 80% to ensure its activity. In addition, we preferentially selected the crRNAs targeting the 20% of the coding region nearest to the start codon. Based on these principles, 10 crRNAs were selected for a given gene. Moreover, the purpose of using this strategy is to attenuate the influence of stochastic factors on the outcomes. If 10 crRNAs could not all be designed within the 20% of the coding region nearest to the start codon, the rest crRNAs were uniformly selected from other regions within the coding region. As a result, a total of 4,153 crRNAs were designed for a crRNA library. Moreover, we also randomly generated 400 negative control crRNAs with no off-target hits in the whole genome based on the above-mentioned criteria.

### Screening experiments

The preparation of the *C. ljungdahlii* competent cells was carried out according to our previously described protocol (Zhao et al., 2019). For the electroporation process, 200 µL of competent cells were mixed with 4 µg of library plasmids and then subject to electroporation (1000V, 50μF, 200Ω) using a Gene Pulser Xcell microbial electroporation system (Bio-Rad, Hercules, CA, USA). With this setting, we could typically obtain around 2.5 × 10^4^ transformants/4µg library plasmids. A total of 10 rounds of electroporations were performed, generating approximately 250,000 transformants, reaching 50-times the size of our crRNA library (4,908 crRNAs). Subsequently, these cells were recovered in 10 mL of the YTF medium at 37°C for 20 h, and then transferred into 600 mL of the YTF medium (containing 5 μg/ml thiamphenicol) and incubated at 37°C. Once the grown cells reached an OD_600_ of ∼ 1.0, 3 µL of the culture was used as the template for PCR amplification to obtain the DNA fragments containing the N24 region of crRNAs, which was used to evaluate the read count distribution of the crRNAs. Meanwhile, this cell library was then subjected to the screening experiments.

To screen for the TF genes capable of affecting the heterotrophic growth of *C. ljungdahlii* (Supplementary Fig. 2), 1.5 mL of the culture from the abovementioned cell library was inoculated into 30 mL of the YTF medium containing 5 μg/ml thiamphenicol. The cell culture was then incubated anaerobically at 37°C with two biological replicates conducted. When the grown cells reached an OD_600_ of ∼ 1.0, 3 µL of the culture was used as the template for PCR amplification to obtain the DNA fragments containing the N24 region of library crRNAs.

To identify the TF genes associated with the autotrophic growth of *C. ljungdahlii* on C1 gases (CO_2_/CO) (Supplementary Fig. 4), 1.5 mL of the culture from the abovementioned cell library was inoculated into 30 mL of the modified ATCC-1754 medium (containing 5 μg/mL of thiamphenicol) with a headspace consisting of syngas (CO-CO_2_-H_2_-N_2_: 56%-20%-9%-15%; pressurized to 0.2 MPa) for gas fermentation. Throughout the fermentation process, the headspace was replenished with fresh gases every 24 h, maintaining a pressure of 0.2 MPa. After 48 h of fermentation, 3 µL of the culture was used as the template for PCR amplification to obtained the DNA fragments containing the N24 region of the library crRNAs.

### NGS library preparation

The N24 regions of the library crRNAs were obtained via PCR amplification using 1 μL of the bacterial solution (OD_600_ = 10) as the template and six pairs of primers with different 8-nt DNA barcodes (TAGGGTGA; TCCTATTC; TTTGGGAA; GTCTGTGC; TGCGACCA and TTGTTGCT) at 5′ends of forward primers to label different PCR products under the following condition: 98°C for 30 s, 22 cycles (98°C for 10 s, 52°C for 30 s, 72°C for 10 s), 72°C for 1 min. The amplification products were resolved on 1.5% agarose gels and the target PCR products were purified using a DNA Gel Extraction Kit (New England Biolab, Ipswich, MA, USA).

The sequencing library was prepared using the VAHTS AmpSeq Library Prep Kit V12 (Vazyme Biotech Co., Ltd., Nanjing, China). Briefly, the fragments of the purified PCR products were treated with End Prep Mix for end repair and 5′ phosphorylation, and then purified using Sample Purification Beads (SPBs). Next, these fragments were adenylated at 3′-ends with A-tailing Mix at 37 °C and ligated to DNA adaptors with a “T” overhang. The products were purified using the SPBs and then subjected to PCR amplification for four cycles using the primers P5/P7. The PCR products were purified with SPBs again, validated using an Agilent 2100 Bioanalyzer (Agilent, Wilmington, DE, USA), and quantified with a Qubit 2.0 Fluorometer (Invitrogen, Carlsbad, CA, USA). Finally, the products with different barcodes were mixed in equal mass and delivered to GENEWIZ for sequencing.

### NGS data processing

The raw NGS data were first de-multiplexed and the adaptor region was removed through Cutadapt to produce clean data. Subsequently, the pairs of paired-end data were merged by FLASH script, and those reads without detected pairs were removed. Python scripts generated in house were then used to search for the ‘AGATN24ATAA’ 32-mer and its corresponding barcode region in the sequencing reads (and the reverse complementary sequence). Note that those carrying mutations within the upstream (AGAT) or downstream (ATAA) flanking regions (4 nt each) were removed. We then mapped the extracted N24 sequences to the designed crRNA library, through which the read count of each crRNA in each library was determined. The read counts were then adjusted using Eq. 1, where *n* is the number of sequencing libraries, to normalize the different sequencing depths of each library.

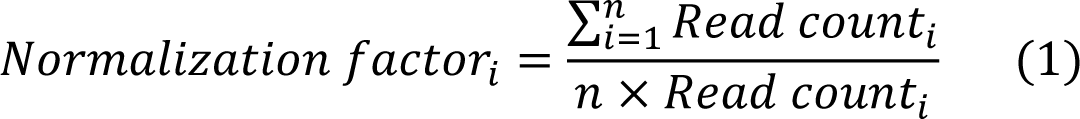

For a given screening condition, the fitness score of each crRNA was calculated with Eq. 2 and Eq. 3. Briefly, we first calculated the abundance variation of each crRNA before and after screening (Eq. 2). Then, each fitness was normalized by subtracting the median fitness of all negative control crRNAs (Eq. 3).

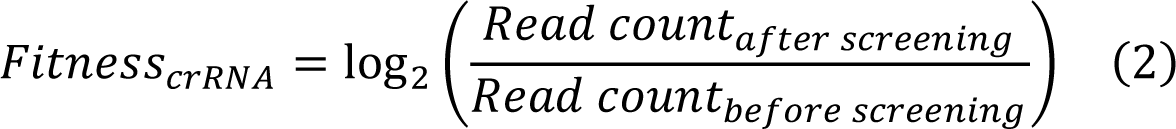

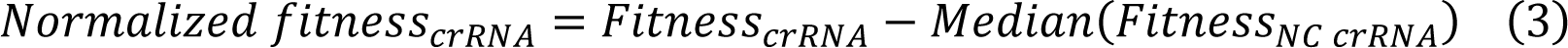

Subsequently, we determined the fitness of each gene and calculated its statistical significance based on the fitness of all crRNAs belonging to this gene. Specifically, the median fitness score of crRNAs of a given gene was defined as the fitness of the gene (Eq. 4).

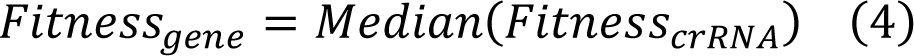

Furthermore, the *P* value for each gene was assessed using a two-tailed Mann-Whitney U (MWU) test, comparing the fitness of its crRNAs against the fitness of all NC crRNAs. The evaluation of the phenotype of each gene was conducted using a pre-defined criterion (Horlbeck et al., 2016) (Eq. 5). This criterion aims to strike a balance between the enrichment of crRNAs in the experiment and its statistical significance. Throughout all experiments, TFs with Score ≥ 0.65 were considered functional.

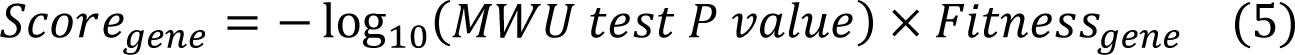

### Overexpression and purification of the His6-tagged NrdR and DeoR protein

The pET28a-*nrdR* and pET28a-*deoR* plasmids were transformed into *E. coli* BL21 (DE3). Gene expression was induced at 16°C for 18 h by adding 0.5 mmol/L Isopropyl-β-D-thio-galactoside (IPTG) when the OD_600_ of the culture reached ∼ 0.8. Cells were harvested by centrifugation (5,000 × *g*, 10 min, 4°C), washed with a solution (pH 7.9, containing 20 mM Tris–HCl, 500 mM KCl, 10% (*v*/*v*) glycerol, and 10 mM imidazole), and then disrupted using a French Press (Constant Systems Limited, Daventry, Northants, UK). Cell debris and membrane fractions were separated from the soluble fraction by centrifugation (20,000 × *g*, 60 min, 4°C). The soluble fraction was loaded onto a Ni Sepharose™ 6 fast flow agarose column (GE Healthcare, Waukesha, WI, USA), and the protein was eluted using a buffer (pH 7.9, containing 20 mM Tris-HCl, 500 mM KCl, 10% (*v*/*v*) glycerol, and 500 mM imidazole). The eluent was then transferred to Amicon Ultra-15 Centrifugal Filter (Milipore, Billerica, MA, USA) and eluted for three times with a buffer (pH 7.9, containing 20 mM Tris-HCl, 500 mM KCl, and 10% (*v*/*v*) glycerol). The purified protein was stored at −80°C.

### Electrophoretic Mobility Shift Assay (EMSA)

DNA probes labeled with cyanine 5 (Cy5) were generated by a two-step PCR amplification process. First, the DNA fragments were obtained by PCR amplification using the *C. ljungdahlii* genomic DNA as the template and specific primer pairs. These primers contained a universal primer sequence (5′-AGCCAGTGGCGATAAG-3′) at the 5′ terminus. Next, a Cy5-tag was incorporated into the abovementioned DNA fragments via PCR amplification using the universal primer labeled with Cy5. The resulting Cy5-labeled probes were subjected to agarose gel electrophoresis and recovered using the AxyPrep™ DNA Gel Extraction Kit (Axygen, Hangzhou, China). EMSAs were performed following the previously described protocol (Ren et al., 2012). In brief, the His6-NrdR protein was pre-incubated with 0.04 pmol Cy5-labeled probes in a buffer (pH 7.9) containing 20 mM Tris-HCl, 5% glycerol, 40 ng/mL bovine serum albumin, 0.25 mM DTT, 10 mM MgCl_2_, 20 mM KCl, and 50 ng/μL fish sperm DNA. The His6-DeoR protein was pre-incubated with 0.04 pmol Cy5-labeled probe in the same buffer except that the pH was adjusted to 11. The mixture was incubated at 25℃ for 20 min. A 7.5% polyacrylamide gel (acrylamide: bis-acrylamide = 80: 1) was prepared and pre-run in 0.5 × TBE buffer at 120 V for 30 min in an ice-bath. Subsequently, 20 μL of the mixture was loaded onto the polyacrylamide gel (20 μl per lane) for electrophoresis (running for 90 min). The gel was visualized using a Starion FLA-9000 Scanner (FujiFilm, Tokyo, Japan).

### Real-time qRT-PCR

*C. ljungdahlii* and its derivative mutants were grown anaerobically. Cells were harvested when OD_600_ reached ∼ 1.0. Total RNA was extracted using RNApure FFPE Kit (CWBIO, Jiangsu, China) and then treated with DNase I (TaKaRa, Kyoto, Japan) to remove DNA contaminants. The concentration of the isolated RNA was determined using a NanoDrop spectrophotometer (Thermo Fisher Scientific, Waltham, MA, USA). To generate cDNA, reverse transcription was performed using the PrimeScript RT reagent kit (TaKaRa, Kyoto, Japan). Real-time qPCR was carried out on a CFX Duet real-time PCR system (Bio-Rad, Hercules, CA, USA) following the steps as previously described (Zhang et al., 2018). The reaction mixtures (20 μl) consisted of 1 × iQ SYBR green Supermix (Bio-Rad, Hercules, CA, USA), 0.5 μM of each primer, and the diluted cDNA template (0.625 ng/μl). For data normalization, the *rho* gene (CLJU_c02220) was adopted as the internal control (Zhao et al., 2019).

### Analytical methods

Cell growth was determined by measuring the absorbance of the culture at *A*_600_ (OD_600_) using a spectrophotometer (DU730, Beckman Coulter, Brea, CA, USA). The concentrations of the products (acetate and ethanol) were measured following the established method (Huang et al., 2016). Briefly, samples were collected at appropriate time intervals and then centrifuged at 7,000 × *g* for 10 min at 4°C. The concentrations of acetate and ethanol in the supernatant were determined using a 7890A gas chromatograph (Agilent, Wilmington, DE, USA) equipped with a capillary column (ECTM-Wax, Alltech, Lexington, KY, USA) and a flame ionization detector (Agilent, Wilmington, DE, USA).

## Acknowledgements

This work was supported by the National Key R&D Program of China (2021YFC2103500), National Natural Science Foundation of China (U2032210 and 31921006), Science and Technology Commission of Shanghai Municipality (21DZ1209100), DNL Cooperation Fund, CAS (DNL202013), and Tianjin Synthetic Biotechnology Innovation Capacity Improvement Project (TSBICIP-KJGG-016).

## Data Availability Statement

The data presented in this study are available on request from the corresponding authors. The raw data supporting these insights for NGS seq have been deposited in the NCBI database with the accession number PRJNA1016157.

