## Supplementary information for "Pooled CRISPR interference screening identifies crucial transcription factors in gas-fermenting *Clostridium ljungdahlii*"

**Supplementary Table 1 Strains and plasmids used in this study**

| **Strains and plasmids** | **Description of genotypes** | **Source** |
| --- | --- | --- |
| **Strains** |  |  |
| ***E. coli*** |  |  |
| BL21 (DE3) | Strain used for protein overexpression | Novagen |
| DH5α | Strain used for genetic manipulation | Invitrogen |
| ***C. ljungdahlii*** |  |  |
| Control | Wild-type *C. ljungdahlii*, carrying the plasmid pZG-ddFncas12a | This study |
| R-c07960 | Wild-type *C. ljungdahlii*, carrying the plasmid pZG-ddFncas12a-c07960 | This study |
| R-c06150 | Wild-type *C. ljungdahlii*, carrying the plasmid pZG-ddFncas12a-c06150 | This study |
| R-c35550 | Wild-type *C. ljungdahlii*, carrying the plasmid pZG-ddFncas12a-c35550 | This study |
| R-c03210 | Wild-type *C. ljungdahlii*, carrying the plasmid pZG-ddFncas12a-c03210 | This study |
| R-c41760 | Wild-type *C. ljungdahlii*, carrying the plasmid pZG-ddFncas12a-c41760 | This study |
| R-c08130 | Wild-type *C. ljungdahlii*, carrying the plasmid pZG-ddFncas12a-c08130 | This study |
| R-c41090 | Wild-type *C. ljungdahlii*, carrying the plasmid pZG-ddFncas12a-c41090 | This study |
| R-c40970 | Wild-type *C. ljungdahlii*, carrying the plasmid pZG-ddFncas12a-c40970 | This study |
| R-c01510 | Wild-type *C. ljungdahlii*, carrying the plasmid pZG-ddFncas12a-c01510 | This study |
| R-*nrdR* | Wild-type *C. ljungdahlii*, carrying the plasmid pZG-ddFncas12a-c12350 | This study |
| R-c11350 | Wild-type *C. ljungdahlii*, carrying the plasmid pZG-ddFncas12a-c11350 | This study |
| R-c42200 | Wild-type *C. ljungdahlii*, carrying the plasmid pZG-ddFncas12a-c42200 | This study |
| R-c24220 | Wild-type *C. ljungdahlii*, carrying the plasmid pZG-ddFncas12a-c24220 | This study |
| R-c41360 | Wild-type *C. ljungdahlii*, carrying the plasmid pZG-ddFncas12a-c41360 | This study |
| R-c39160 | Wild-type *C. ljungdahlii*, carrying the plasmid pZG-ddFncas12a-c39160 | This study |
| R-c12190 | Wild-type *C. ljungdahlii*, carrying the plasmid pZG-ddFncas12a-c12190 | This study |
| R-c08220 | Wild-type *C. ljungdahlii*, carrying the plasmid pZG-ddFncas12a-c08220 | This study |
| R-c01230 | Wild-type *C. ljungdahlii*, carrying the plasmid pZG-ddFncas12a-c01230 | This study |
| R-c31510 | Wild-type *C. ljungdahlii*, carrying the plasmid pZG-ddFncas12a-c31510 | This study |
| R-c11190 | Wild-type *C. ljungdahlii*, carrying the plasmid pZG-ddFncas12a-c11190 | This study |
| R-c30510 | Wild-type *C. ljungdahlii*, carrying the plasmid pZG-ddFncas12a-c30510 | This study |
| R-c01560 | Wild-type *C. ljungdahlii*, carrying the plasmid pZG-ddFncas12a-c01560 | This study |
| R-c12100 | Wild-type *C. ljungdahlii*, carrying the plasmid pZG-ddFncas12a-c12100 | This study |
| WT | Wild-type *C. ljungdahlii*, DSM 13528 | DSMZ |
| △c12350 | Wild-type *C. ljungdahlii* with the deletion of *nrdR* (CLJU_c12350) | This study |
| WT (P) | Wild-type *C. ljungdahlii*, carrying the plasmid pMTL-c83151 | This study |
| P*_1339_*-c07960 | Wild-type *C. ljungdahlii*, carrying the plasmid pMTL83151- P*_1339_*-c07960 | This study |
| P*_1339_*-c20610 | Wild-type *C. ljungdahlii*, carrying the plasmid pMTL83151- P*_1339_*-c20610 (*deoR*) | This study |
| P*_1339_*-c34000 | Wild-type *C. ljungdahlii*, carrying the plasmid pMTL83151- P*_1339_*-c34000 | This study |
| P*_pta_*-*deoR* | Wild-type *C. ljungdahlii*, carrying the plasmid pMTL83151- P*_pta_*-*deoR* | This study |
| P*_ptb_*-*deoR* | Wild-type *C. ljungdahlii*, carrying the plasmid pMTL83151- P*_ptb_*-*deoR* | This study |
| P*_01440_*-*deoR* | Wild-type *C. ljungdahlii*, carrying the plasmid pMTL83151- P*_01440_*-*deoR* | This study |
| P*_thl_*- *deoR* | Wild-type *C. ljungdahlii*, carrying the plasmid pMTL83151- P*_thl_*- *deoR* | This study |
| P*_1339_*-c11350 | Wild-type *C. ljungdahlii*, carrying the plasmid pMTL83151- P*_1339_*-c11350 | This study |
| P*_1339_*-c42200 | Wild-type *C. ljungdahlii*, carrying the plasmid pMTL83151- P*_1339_*-c42200 | This study |
| P*_1339_*-c41360 | Wild-type *C. ljungdahlii*, carrying the plasmid pMTL83151- P*_1339_*-c41360 | This study |
| P*_1339_*-c31510 | Wild-type *C. ljungdahlii*, carrying the plasmid pMTL83151- P*_1339_*-c31510 | This study |
| P*_1339_*-c14130 | Wild-type *C. ljungdahlii*, carrying the plasmid pMTL83151- P*_1339_*-c14130 | This study |
| P*_1339_*-c34650 | Wild-type *C. ljungdahlii*, carrying the plasmid pMTL83151- P*_1339_*-c34650 | This study |
| P*_1339_*-c12100 | Wild-type *C. ljungdahlii*, carrying the plasmid pMTL83151- P*_1339_*-c12100 | This study |
| P*_1339_*-c35550 | Wild-type *C. ljungdahlii*, carrying the plasmid pMTL83151- P*_1339_*-c35550 | This study |
| P*_1339_*-c30510 | Wild-type *C. ljungdahlii*, carrying the plasmid pMTL83151- P*_1339_*-c30510 | This study |
| P*_1339_*-c28220 | Wild-type *C. ljungdahlii*, carrying the plasmid pMTL83151- P*_1339_*-c28220 | This study |
| P*_1339_*-c32060 | Wild-type *C. ljungdahlii*, carrying the plasmid pMTL83151- P*_1339_*-c32060 | This study |
| P*_1339_*-c01860 | Wild-type *C. ljungdahlii*, carrying the plasmid pMTL83151- P*_1339_*-c01860 | This study |
| P*_1339_*-c37800 | Wild-type *C. ljungdahlii*, carrying the plasmid pMTL83151- P*_1339_*-c37800 | This study |
| **Plasmids** |  |  |
| pZG-ddFncas12a | *ColE1, catP, pCB102 ori, P_01440_-Fncas12a, P_thl_* | This lab (Zhao et al., 2019) |
| pZG-ddFncas12a-c07960 | Repression vector targeting CLJU_c07960 | This study |
| pZG-ddFncas12a-c06150 | Repression vector targeting CLJU_c06150 | This study |
| pZG-ddFncas12a-c35550 | Repression vector targeting CLJU_c35550 | This study |
| pZG-ddFncas12a-c03210 | Repression vector targeting CLJU_c03210 | This study |
| pZG-ddFncas12a-c41760 | Repression vector targeting CLJU_c41760 | This study |
| pZG-ddFncas12a-c08130 | Repression vector targeting CLJU_c08130 | This study |
| pZG-ddFncas12a-c41090 | Repression vector targeting CLJU_c41090 | This study |
| pZG-ddFncas12a-c40970 | Repression vector targeting CLJU_c40970 | This study |
| pZG-ddFncas12a-c01510 | Repression vector targeting CLJU_c01510 | This study |
| pZG-ddFncas12a-c12350 | Repression vector targeting CLJU_c12350 | This study |
| pZG-ddFncas12a-c11350 | Repression vector targeting CLJU_c11350 | This study |
| pZG-ddFncas12a-c42200 | Repression vector targeting CLJU_c42200 | This study |
| pZG-ddFncas12a-c24220 | Repression vector targeting CLJU_c24220 | This study |
| pZG-ddFncas12a-c41360 | Repression vector targeting CLJU_c41360 | This study |
| pZG-ddFncas12a-c39160 | Repression vector targeting CLJU_c39160 | This study |
| pZG-ddFncas12a-c12190 | Repression vector targeting CLJU_c12190 | This study |
| pZG-ddFncas12a-c08220 | Repression vector targeting CLJU_c08220 | This study |
| pZG-ddFncas12a-c01230 | Repression vector targeting CLJU_c01230 | This study |
| pZG-ddFncas12a-c31510 | Repression vector targeting CLJU_c31510 | This study |
| pZG-ddFncas12a-c11190 | Repression vector targeting CLJU_c11190 | This study |
| pZG-ddFncas12a-c30510 | Repression vector targeting CLJU_c30510 | This study |
| pZG-ddFncas12a-c01560 | Repression vector targeting CLJU_c01560 | This study |
| pZG-ddFncas12a-c12100 | Repression vector targeting CLJU_c12100 | This study |
| pMTLcas-*pta* | *pCB102 ori, catP, ColE1,tra, P_thl_-Cas9, P_araE_-sgRNA, pta homologous arm* | This lab (Huang et al., 2016) |
| pMTLcas-*nrdR* | *ColE1, catP, pCB102 ori, P_thl_-Cas9, P_1339_-sgRNA, nrdR homologous arm* | This study |
| pMTL83151 | *ColE1, catP, pCB102 ori* | Provided by Prof. Nigel P. Minton (Heap et al., 2009) |
| pMTL83151- P*_1339_*-c07960 | Vector for the overexpression of CLJU_c07960 with the promoter P*_1339_* | This study |
| pMTL83151- P*_1339_*-c20610 | Vector for the overexpression of CLJU_c20610 with the promoter P*_1339_* | This study |
| pMTL83151- P*_1339_*-c34000 | Vector for the overexpression of CLJU_c34000 with the promoter P*_1339_* | This study |
| pMTL83151- P*_pta_*-*deoR* | Vector for the overexpression of *deoR* with the promoter P*_pta_* | This study |
| pMTL83151- P*_ptb_*-*deoR* | Vector for the overexpression of *deoR* with the promoter P*_ptb_* | This study |
| pMTL83151- P*_01440_*-*deoR* | Vector for the overexpression of *deoR* with the promoter P*_01440_* | This study |
| pMTL83151- P*_thl_*- *deoR* | Vector for the overexpression of *deoR* with the promoter P*_thl_* | This study |
| pMTL83151- P*_1339_*-c11350 | Vector for the overexpression of CLJU_c11350 with the promoter P*_1339_* | This study |
| pMTL83151- P*_1339_*-c42200 | Vector for the overexpression of CLJU_c42200 with the promoter P*_1339_* | This study |
| pMTL83151- P*_1339_*-c41360 | Vector for the overexpression of CLJU_c41360 with the promoter P*_1339_* | This study |
| pMTL83151- P*_1339_*-c31510 | Vector for the overexpression of CLJU_c31500 with the promoter P*_1339_* | This study |
| pMTL83151- P*_1339_*-c14130 | Vector for the overexpression of CLJU_c14130 with the promoter P*_1339_* | This study |
| pMTL83151- P*_1339_*-c34650 | Vector for the overexpression of CLJU_c34650 with the promoter P*_1339_* | This study |
| pMTL83151- P*_1339_*-c12100 | Vector for the overexpression of CLJU_c12100 with the promoter P*_1339_* | This study |
| pMTL83151- P*_1339_*-c35550 | Vector for the overexpression of CLJU_c35550 with the promoter P*_1339_* | This study |
| pMTL83151- P*_1339_*-c30510 | Vector for the overexpression of CLJU_c34510 with the promoter P*_1339_* | This study |
| pMTL83151- P*_thl_*-c28220 | Vector for the overexpression of CLJU_c28220 with the promoter P*_thl_* | This study |
| pMTL83151- P*_thl_*-c32060 | Vector for the overexpression of CLJU_c32060 with the promoter P*_thl_* | This study |
| pMTL83151- P*_thl_*-c01860 | Vector for the overexpression of CLJU_c01860 with the promoter P*_thl_* | This study |
| pMTL83151- P*_thl_*-c37800 | Vector for the overexpression of CLJU_c37800 with the promoter P*_thl_* | This study |

**Supplementary Table 2 Primers used in this study**

| **Primer name** | **Sequence (5′‒3′)** | **Description** |
| --- | --- | --- |
| Universal-for | CCCCGTATCAAAATTTAGGAGGTTAGGATCCAATTTCTACTGTTGTAGAT | Universal forward primer for gene repression |
| Universal-rev | CCGTCGACCCCGGGCCATGGATAAAAATAAGAAGCCTGCAAATGCAGG | Universal reverse primer for gene repression |
| c07960-crRNA-for | GGATCCAATTTCTACTGTTGTAGATGAACAACTTCACAGTTCTTCAGGA | Forward primer used for constructing crRNA targeting CLJU_c07960 |
| c07960-crRNA-rev | TAAGAAGCCTGCAAATGCAGGCTTCTTATTTTTATTCCTGAAGAACTGTGAAGTTGTTC | Reverse primer used for constructing crRNA targeting CLJU_c07960 |
| c06150-crRNA-for | GGATCCAATTTCTACTGTTGTAGATAATACTTCCATTGAGTATTTAATG | Forward primer used for constructing crRNA targeting CLJU_c06150 |
| c06150-crRNA-rev | TAAGAAGCCTGCAAATGCAGGCTTCTTATTTTTATCATTAAATACTCAATGGAAGTATT | Reverse primer used for constructing crRNA targeting CLJU_c06150 |
| c35550-crRNA-for | GGATCCAATTTCTACTGTTGTAGATCAATCTACCAAGAAGCATCCTTCG | Forward primer used for constructing crRNA targeting CLJU_c35550 |
| c35550-crRNA-rev | TAAGAAGCCTGCAAATGCAGGCTTCTTATTTTTATCGAAGGATGCTTCTTGGTAGATTG | Reverse primer used for constructing crRNA targeting CLJU_c35550 |
| c03210-crRNA-for | GGATCCAATTTCTACTGTTGTAGATCTGAAAATGCTGCAATTGCCAGAT | Forward primer used for constructing crRNA targeting CLJU_c03210 |
| c03210-crRNA-rev | TAAGAAGCCTGCAAATGCAGGCTTCTTATTTTTATATCTGGCAATTGCAGCATTTTCAG | Reverse primer used for constructing crRNA targeting CLJU_c03210 |
| c41760-crRNA-for | GGATCCAATTTCTACTGTTGTAGATATGATGGTAAGACAGCGGAAACTA | Forward primer used for constructing crRNA targeting CLJU_c41760 |
| c41760-crRNA-rev | TAAGAAGCCTGCAAATGCAGGCTTCTTATTTTTATTAGTTTCCGCTGTCTTACCATCAT | Reverse primer used for constructing crRNA targeting CLJU_c41760 |
| c08130-crRNA-for | GGATCCAATTTCTACTGTTGTAGATAGACCTGATTTAGCCATACTTACA | Forward primer used for constructing crRNA targeting CLJU_c08130 |
| c08130-crRNA-rev | TAAGAAGCCTGCAAATGCAGGCTTCTTATTTTTATTGTAAGTATGGCTAAATCAGGTCT | Reverse primer used for constructing crRNA targeting CLJU_c08130 |
| c41090-crRNA-for | GGATCCAATTTCTACTGTTGTAGATACAAATGGATATGAAGTAAGTGCG | Forward primer used for constructing crRNA targeting CLJU_c41090 |
| c41090-crRNA-rev | TAAGAAGCCTGCAAATGCAGGCTTCTTATTTTTATCGCACTTACTTCATATCCATTTGT | Reverse primer used for constructing crRNA targeting CLJU_c41090 |
| c40970-crRNA-for | GGATCCAATTTCTACTGTTGTAGATGGTAGCAGTTGTTAAACCAGGTAG | Forward primer used for constructing crRNA targeting CLJU_c40970 |
| c40970-crRNA-rev | TAAGAAGCCTGCAAATGCAGGCTTCTTATTTTTATCTACCTGGTTTAACAACTGCTACC | Reverse primer used for constructing crRNA targeting CLJU_c40970 |
| c01510-crRNA-for | GGATCCAATTTCTACTGTTGTAGATTGCGGTGACGCAAGAGACGTTGTA | Forward primer used for constructing crRNA targeting CLJU_c01510 |
| c01510-crRNA-rev | TAAGAAGCCTGCAAATGCAGGCTTCTTATTTTTATTACAACGTCTCTTGCGTCACCGCA | Reverse primer used for constructing crRNA targeting CLJU_c01510 |
| *nrdR*-crRNA-for | GGATCCAATTTCTACTGTTGTAGATAGGTGTAATAAAAGATATACCACT | Forward primer used for constructing crRNA targeting CLJU_c12350 |
| *nrdR*-crRNA-rev | TAAGAAGCCTGCAAATGCAGGCTTCTTATTTTTATAGTGGTATATCTTTTATTACACCT | Reverse primer used for constructing crRNA targeting CLJU_c12350 |
| c11350-crRNA-for | GGATCCAATTTCTACTGTTGTAGATTGGTGCTTGTCCTGGTGATAATGC | Forward primer used for constructing crRNA targeting CLJU_c11350 |
| c11350-crRNA-rev | TAAGAAGCCTGCAAATGCAGGCTTCTTATTTTTATGCATTATCACCAGGACAAGCACCA | Reverse primer used for constructing crRNA targeting CLJU_c11350 |
| c42200-crRNA-for | TAAGAAGCCTGCAAATGCAGGCTTCTTATTTTTATGCATTATCACCAGGACAAGCACCA | Forward primer used for constructing crRNA targeting CLJU_c42200 |
| c42200-crRNA-rev | TAAGAAGCCTGCAAATGCAGGCTTCTTATTTTTATTCGTGTAACATATTGCTTTCTTTC | Reverse primer used for constructing crRNA targeting CLJU_c42200 |
| c24220-crRNA-for | GGATCCAATTTCTACTGTTGTAGATCCGAGTTGTCTGATATGTATTATG | Forward primer used for constructing crRNA targeting CLJU_c24220 |
| c24220-crRNA-rev | TAAGAAGCCTGCAAATGCAGGCTTCTTATTTTTATCATAATACATATCAGACAACTCGG | Reverse primer used for constructing crRNA targeting CLJU_c24220 |
| c41360-crRNA-for | GGATCCAATTTCTACTGTTGTAGATAAATTGGCAGAAACGAACTAGCTA | Forward primer used for constructing crRNA targeting CLJU_c41360 |
| c41360-crRNA-rev | TAAGAAGCCTGCAAATGCAGGCTTCTTATTTTTATTAGCTAGTTCGTTTCTGCCAATTT | Reverse primer used for constructing crRNA targeting CLJU_c41360 |
| c39160-crRNA-for | GGATCCAATTTCTACTGTTGTAGATGCCAATAACCTAGGTATAGGTGAG | Forward primer used for constructing crRNA targeting CLJU_c39160 |
| c39160-crRNA-rev | TAAGAAGCCTGCAAATGCAGGCTTCTTATTTTTATCTCACCTATACCTAGGTTATTGGC | Reverse primer used for constructing crRNA targeting CLJU_c39160 |
| c12190-crRNA-for | GGATCCAATTTCTACTGTTGTAGATCGCCGCAGAGGCGTGCTGTAGTTG | Forward primer used for constructing crRNA targeting CLJU_c12190 |
| c12190-crRNA-rev | TAAGAAGCCTGCAAATGCAGGCTTCTTATTTTTATCAACTACAGCACGCCTCTGCGGCG | Reverse primer used for constructing crRNA targeting CLJU_c12190 |
| c08220-crRNA-for | GGATCCAATTTCTACTGTTGTAGATGATTTATCGGTTCCTGATGGAATT | Forward primer used for constructing crRNA targeting CLJU_c08220 |
| c08220-crRNA-rev | TAAGAAGCCTGCAAATGCAGGCTTCTTATTTTTATAATTCCATCAGGAACCGATAAATC | Reverse primer used for constructing crRNA targeting CLJU_c08220 |
| c01230-crRNA-for | TCCAATTTCTACTGTTGTAGATCAACAGAATCGGAACTTATGGAGC | Forward primer used for constructing crRNA targeting CLJU_c01230 |
| c01230-crRNA-rev | TAAGAAGCCTGCAAATGCAGGCTTCTTATTTTTATGCTCCATAAGTTCCGATTCTGTTG | Reverse primer used for constructing crRNA targeting CLJU_c01230 |
| c31510-crRNA-for | GGATCCAATTTCTACTGTTGTAGATATGGAGAAAGGCTCAAGTCTGCCA | Forward primer used for constructing crRNA targeting CLJU_c31510 |
| c31510-crRNA-rev | TAAGAAGCCTGCAAATGCAGGCTTCTTATTTTTATTGGCAGACTTGAGCCTTTCTCCAT | Reverse primer used for constructing crRNA targeting CLJU_c31510 |
| c11190-crRNA-for | GGATCCAATTTCTACTGTTGTAGATGCGGATAGACTAAAAAAAAGCGGT | Forward primer used for constructing crRNA targeting CLJU_c11190 |
| c11190-crRNA-rev | TAAGAAGCCTGCAAATGCAGGCTTCTTATTTTTATACCGCTTTTTTTTAGTCTATCCGC | Reverse primer used for constructing crRNA targeting CLJU_c11190 |
| c30510-crRNA-for | GGATCCAATTTCTACTGTTGTAGATATGGTAATCTTGAACCAGGAGATA | Forward primer used for constructing crRNA targeting CLJU_c30510 |
| c30510-crRNA-rev | TAAGAAGCCTGCAAATGCAGGCTTCTTATTTTTATTATCTCCTGGTTCAAGATTACCAT | Reverse primer used for constructing crRNA targeting CLJU_c30510 |
| c01560-crRNA-for | GGATCCAATTTCTACTGTTGTAGATAGTTATATGAGACCTATATTAACC | Forward primer used for constructing crRNA targeting CLJU_c01560 |
| c01560-crRNA-rev | TAAGAAGCCTGCAAATGCAGGCTTCTTATTTTTATGGTTAATATAGGTCTCATATAACT | Reverse primer used for constructing crRNA targeting CLJU_c01560 |
| c12100-crRNA-for | GGATCCAATTTCTACTGTTGTAGATGCTATTCACTATAGTGATGAGCCG | Forward primer used for constructing crRNA targeting CLJU_c12100 |
| c12100-crRNA-rev | TAAGAAGCCTGCAAATGCAGGCTTCTTATTTTTATCGGCTCATCACTATAGTGAATAGC | Reverse primer used for constructing crRNA targeting CLJU_c12100 |
| *nrdR*-sgRNA-for | CTTAAGGAGGAGTTTTCGTCGACAATAATTTCCCCTATCTCCGGTTTTAGAGCTAGAAA | Forward primer of *nrdR* sgRNA |
| sgRNA-rev | ATAAAAATAAGAAGCCTGCAAATGCAGGCTTCTTATTTTTATAAAAAAAGCACCGACTC | Universal reverse primer of sgRNA |
| *nrdR*-UpArm-for | AGCCTGCATTTGCAGGCTTCTTATTTTTATATGATAATTAATAAAGTAGAAATTTG | Forward primer of *nrdR*’s up homologous arm |
| *nrdR*-UpArm-rev | ACCATTTATTTATTTGAAATAAGTTTAATTCACGTCCTTCATATAAGATAG | Reverse primer of *nrdR*’s up homologous arm |
| *nrdR*-DownArm-for | CTATCTTATATGAAGGACGTGAATTAAACTTATTTCAAATAAATAAATGGT | Forward primer of *nrdR*’s down homologous arm |
| *nrdR*-DownArm-rev | CTTGCATGTCTGCAGGCCTCGAGGCACACATCATATCCATCCA | Reverse primer of nrdr’s down homologous arm |
| P*_1339_*-for | GCGGCCGCTGTATCCATATGTTTATATTTAGTCCCTTGCCTTG | Forward primer used for amplifying P*_1339_* |
| P*_1339_*-rev | GAAAACTCCTCCTTAAGATTTATATATGTG | Reverse primer used for amplifying P*_1339_* |
| G*_07960_* (P*_1339_*)-for | CACATATATAAATCTTAAGGAGGAGTTTTCATGGAAATGGATGAAAGAAA | Forward primer used for amplifying CLJU_c07960 |
| G*_07960_*-rev | ACGGCCAGTGCCAAGCTTCTATTTTTCATCAGAATAGGCA | Reverse primer used for amplifying CLJU_c07960 |
| *deoR* (P*_1339_*)-for | CACATATATAAATCTTAAGGAGGAGTTTTCTTGCTAACAGAACAGCGCCA | Forward primer used for amplifying *deoR* |
| G*_34000_* (P*_1339_*)-for | CACATATATAAATCTTAAGGAGGAGTTTTCATGAAAATTACACAGGAATCA | Forward primer used for amplifying CLJU_c34000 |
| G*_34000_*-rev | ACGGCCAGTGCCAAGCTTTTAAATTTTATCTTCCATTATATCTT | Reverse primer used for amplifying CLJU_c34000 |
| P*_pta_*-for | GCGGCCGCTGTATCCATATGAGAAATTTTCCTTTCTAAAATATTT | Forward primer used for amplifying P*_pta_* |
| P*_pta_*-rev | GTTCATTTCCTCCCTTTAAA | Reverse primer used for amplifying P*_pta_* |
| *deoR* (P*_pta_*)-for | TTTAAAGGGAGGAAATGAACTTGCTAACAGAACAGCGCCA | Forward primer used for amplifying *deoR* |
| P*_ptb_*-for | GCGGCCGCTGTATCCATATGTATAAAATATAAATAATTTTCTAAAAAACTTAAC | Forward primer used for amplifying P*_ptb_* |
| P*_ptb_*-rev | TCGACACTCCCTTTTACTA | Reverse primer used for amplifying P*_ptb_* |
| *deoR* (P*_ptb_*)-for | TAGTAAAAGGGAGTGTCGATTGCTAACAGAACAGCGCCA | Forward primer used for amplifying *deoR* |
| P*_01440_*-for | GCGGCCGCTGTATCCATATGGGCATTTTCAAAGAAATAACTAG | Forward primer used for amplifying P*_01440_* |
| P*_01440_*-rev | CTTATGTAACACCTCCTTAATTTTTAG | Forward primer used for amplifying P*_01440_* |
| *deoR* (P*_01440_*)-for | CTAAAAATTAAGGAGGTGTTACATAAGTTGCTAACAGAACAGCGCCA | Forward primer used for amplifying *deoR* |
| P*_thl_*-for | GCGGCCGCTGTATCCATATGTTTTTAACAAAATATATTGATAAAAATAAT | Forward primer used for amplifying P*_thl_* |
| P*_thl_*-rev | TCTAACTAACCTCCTAAAT | Forward primer used for amplifying P*_thl_* |
| *deoR* (P*_thl_*)-for | ATTTAGGAGGTTAGTTAGATTGCTAACAGAACAGCGCCA | Forward primer used for amplifying *deoR* |
| *deoR*-rev | ACGGCCAGTGCCAAGCTTTTAATCTGTCACAACCTTTACT | Reverse primer used for amplifying *deoR* |
| G*_11350_* (P*_1339_*)-for | CACATATATAAATCTTAAGGAGGAGTTTTCATGAAAAGAGAATCGGAGG | Forward primer used for amplifying CLJU_c11350 |
| G*_11350_*-rev | ACGGCCAGTGCCAAGCTTTCAATACAATATCTTTGTGATTACT | Reverse primer used for amplifying CLJU_c11350 |
| G*_42200_* (P*_1339_*)-for | CACATATATAAATCTTAAGGAGGAGTTTTCGTGAACAAAGGTATAAATGTTAT | Forward primer used for amplifying CLJU_c42200 |
| G*_422000_*-rev | ACGGCCAGTGCCAAGCTTTTATTCATTATTCTTCTCCTTTG | Reverse primer used for amplifying CLJU_c42200 |
| G*_41360_* (P*_1339_*)-for | CACATATATAAATCTTAAGGAGGAGTTTTCTTGGCAAGATTATCAGATAT | Forward primer used for amplifying CLJU_c41360 |
| G*_41360_*-rev | ACGGCCAGTGCCAAGCTTCTATCTATTCATACCAAATTTAGG | Reverse primer used for amplifying CLJU_c41360 |
| G*_31510_* (P*_1339_*)-for | CACATATATAAATCTTAAGGAGGAGTTTTCATGTTAAATGGTAATAAAGAGGATGTAGG | Forward primer used for amplifying CLJU_c31510 |
| G*_31510_*-rev | ACGGCCAGTGCCAAGCTTTTATTTATAGCTAAATTGTTCAGTATTTTTTATCG | Reverse primer used for amplifying CLJU_c31510 |
| G*_14130_* (P*_1339_*)-for | CACATATATAAATCTTAAGGAGGAGTTTTCATGAACAGGAATAAAGCGGA | Forward primer used for amplifying CLJU_c14130 |
| G*_14130_*-rev | ACGGCCAGTGCCAAGCTTCTATTGAAAATCATTTTGATCTT | Reverse primer used for amplifying CLJU_c14130 |
| G*_34650_* (P*_1339_*)-for | CACATATATAAATCTTAAGGAGGAGTTTTCATGTTTGAAAATACTTTAGAACT | Forward primer used for amplifying CLJU_c34650 |
| G*_34650_*-rev | ACGGCCAGTGCCAAGCTTTTAATCTTCCATAAGTAATTCAAC | Reverse primer used for amplifying CLJU_c34650 |
| G*_12100_* (P*_1339_*)-for | CACATATATAAATCTTAAGGAGGAGTTTTCTTGAAGCTATCCACAAAAGG | Forward primer used for amplifying CLJU_c12100 |
| G*_12100_*-rev | ACGGCCAGTGCCAAGCTTTTATATATCCCCTTTCGTTTTT | Reverse primer used for amplifying CLJU_c12100 |
| G*_35550_* (P*_1339_*)-for | CACATATATAAATCTTAAGGAGGAGTTTTCATGGATAATTTAACTTCTATTTTTAG | Forward primer used for amplifying CLJU_c35550 |
| G*_35550_*-rev | ACGGCCAGTGCCAAGCTTTTATTCTTTAGAATTTTTTTGGCA | Reverse primer used for amplifying CLJU_c35550 |
| G*_30510_* (P*_1339_*)-for | CACATATATAAATCTTAAGGAGGAGTTTTCATGGGTTCACTATTTGAAAAAATTGAAG | Forward primer used for amplifying CLJU_c30510 |
| G*_30510_*-rev | ACGGCCAGTGCCAAGCTTTTAAAATAAATGAACTTCTTTTTCAATAAACTTC | Reverse primer used for amplifying CLJU_c30510 |
| G*_28220_* (P*_thl_*)-for | CAAAATTTAGGAGGTTAGTTAGAATGGACATTAGCACCAAATT | Forward primer used for amplifying CLJU_c28220 |
| G*_28220_*-rev | ACGGCCAGTGCCAAGCTTCTAATTTTTATTAGTTTCAAAATTAAT | Reverse primer used for amplifying CLJU_c28220 |
| G*_32060_* (P*_thl_*)-for | CAAAATTTAGGAGGTTAGTTAGAATGTTTTCGGAACTTAAGAT | Forward primer used for amplifying CLJU_c32060 |
| G*_32060_*-rev | ACGGCCAGTGCCAAGCTTTCAAGTCCATCCTCCTATT | Reverse primer used for amplifying CLJU_c32060 |
| G*_01860_* (P*_thl_*)-for | CAAAATTTAGGAGGTTAGTTAGAATGAATAACAATTCGGAAATTG | Forward primer used for amplifying CLJU_c01860 |
| G*_01860_*-rev | ACGGCCAGTGCCAAGCTTCTAATTATGTTTTACATGTTCTAATC | Reverse primer used for amplifying CLJU_c01860 |
| G*_37800_* (P*_thl_*)-for | CAAAATTTAGGAGGTTAGTTAGAATGACTTTAAGACATTTTAAAATAT | Forward primer used for amplifying CLJU_c37800 |
| G*_37800_*-rev | ACGGCCAGTGCCAAGCTTTTACTTCTTTATTGTACATAAATC | Reverse primer used for amplifying CLJU_c37800 |
| qRT-*nrdD*-for | AGGCTATGTTAAAATTGGGC | Forward qrt-PCR primer for *nrdD* |
| qRT-*nrdD*-rev | AAAAGCATTTCCTTTGTTGG | Reverse qrt-PCR primer for *nrdD* |
| qRT-*nrdA*-for | AGCAGAAGCCTGAGTCCTTG | Forward qrt-PCR primer for *nrdA* |
| qRT-*nrdA*-rev | GGAGTTCCCGGAGAAAATGG | Reverse qrt-PCR primer for *nrdA* |
| qRT-c20040-for | CGCGTTCTTTAGCTTTTAGC | Forward qrt-PCR primer for CLJU_c20040 |
| qRT-c20040-rev | AGGAAGCAATTGGCTTTATT | Reverse qrt-PCR primer for CLJU_c20040 |
| qRT-c17910-for | GCCCCTGCAGAATTAATCTT | Forward qrt-PCR primer for CLJU_c17910 |
| qRT-c17910-rev | ACACTGGGTTCCTCTACATC | Reverse qrt-PCR primer for CLJU_c17910 |
| qRT-c37670-for | GGTGGTAAATTAACTTCTCC | Forward qrt-PCR primer for CLJU_c37670 |
| qRT-c37670-rev | GGAAAAGGTGTAGAAAGAGG | Reverse qrt-PCR primer for CLJU_c37670 |
| qRT-c16510-for | AGAGAGAGATGCAGGATTTG | Forward qrt-PCR primer for CLJU_c16510 |
| qRT-c16510-rev | AATTTGCTTCTCCCATTACC | Reverse qrt-PCR primer for CLJU_c16510 |
| EMSA-*nrdD*-for | AGCCAGTGGCGATAAGATTTTGCCGATCATGAAAGT | Forward primer for EMSA |
| EMSA*-nrdD*-rev | AGCCAGTGGCGATAAGCCATCTCTTTTCACAACATG | Reverse primer for EMSA |
| EMSA-*nrdA*-for | AGCCAGTGGCGATAAGAACAAGTCTTTTACGCAGTC | Forward primer for EMSA |
| EMSA*-nrdA*-rev | AGCCAGTGGCGATAAGCTCCTTAAGAATTTAATATTTGAAAACC | Reverse primer for EMSA |
| EMSA-c06920-for | AGCCAGTGGCGATAAGCCTCTCATTATGTCTAATTCTG | Forward primer for EMSA |
| EMSA-c06920-rev | AGCCAGTGGCGATAAGCTGCTACATTTGTTAAAGGAC | Reverse primer for EMSA |
| EMSA-c20040-for | AGCCAGTGGCGATAAGGGACAAGTAGTTAGTATACTTTTC | Forward primer for EMSA |
| EMSA-c20040-rev | AGCCAGTGGCGATAAGCTTTATATCTTTGAAAAGTGTTCG | Reverse primer for EMSA |
| EMSA-c17910-for | AGCCAGTGGCGATAAGGCTCTGAAAGCAGAAGAATT | Forward primer for EMSA |
| EMSA-c17910-rev | AGCCAGTGGCGATAAGCTGTTCTTTCTCGTATAGTTTC | Reverse primer for EMSA |
| EMSA-c37670-for | AGCCAGTGGCGATAAGCCTGATCAATTGATTTTGCTT | Forward primer for EMSA |
| EMSA-c37670-rev | AGCCAGTGGCGATAAGCGCAATTGCATATTTCAAACA | Reverse primer for EMSA |
| EMSA-c16510-for | AGCCAGTGGCGATAAGCTTTAGTTGGTTTATCATGAT | Forward primer for EMSA |
| EMSA-c16510-rev | AGCCAGTGGCGATAAGTTATGCATAATTATTTATATATTGTC | Reverse primer for EMSA |
| EMSA-c37630-for | AGCCAGTGGCGATAAGCCTGTTATAGCCATTTTCAT | Forward primer for EMSA |
| EMSA-c37630-rev | AGCCAGTGGCGATAAGGAATAAAGACAAGCAATTGCC | Reverse primer for EMSA |

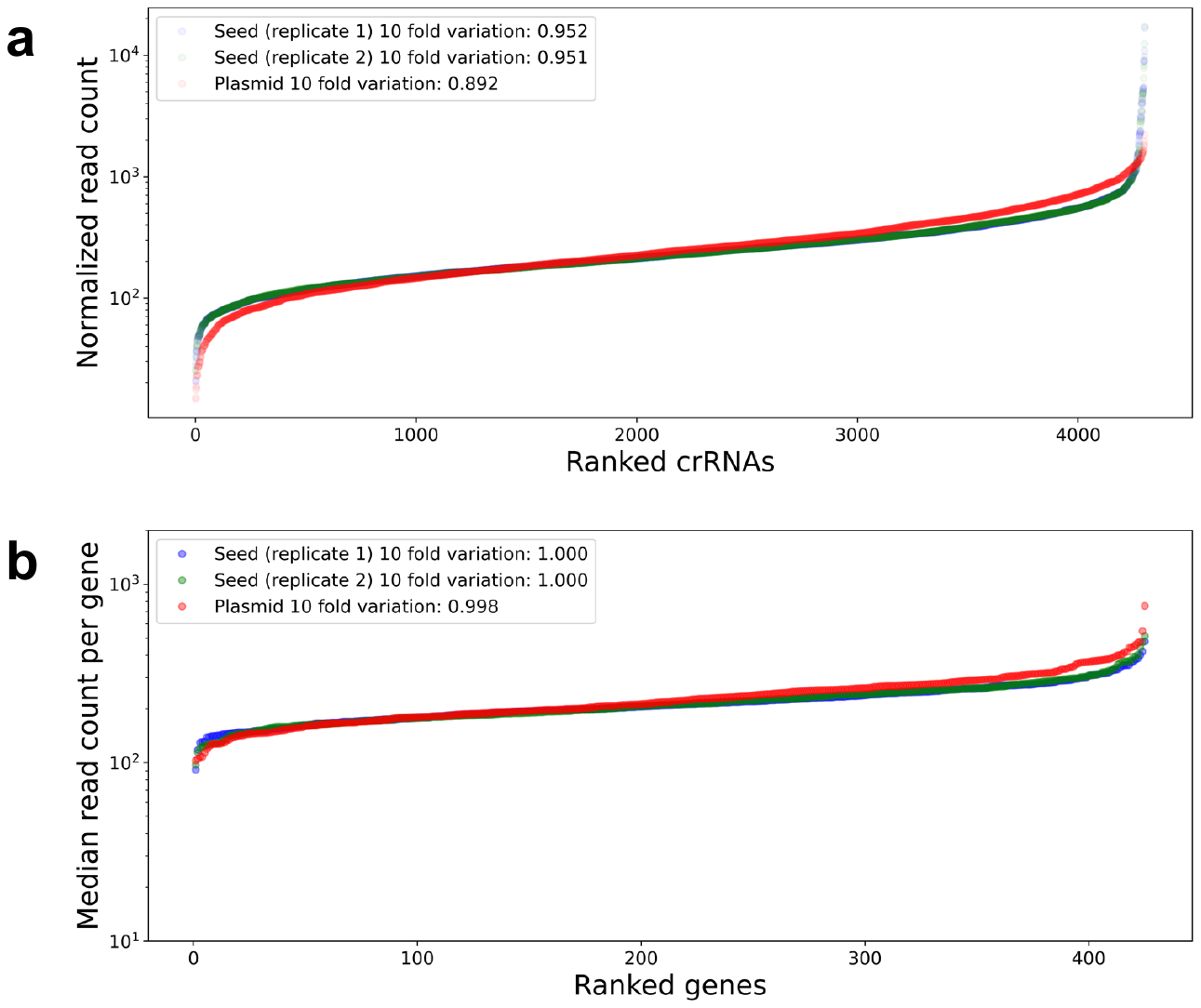

**Supplementary Fig. 1** NGS profiling of crRNA library. **a**, Distribution of crRNA read counts in ascending order. **b**, Distribution of median crRNA read counts of each gene in ascending order. In **a** and **b**, red, purple, and green dots represent the plasmid library, replicate 1 of seed culture, and replicate 2 of seed culture, respectively. A 10-fold variation is displayed for all distribution profiles, which were computed by summarizing the event count (crRNA or gene) with read counts falling within the range of [1/3.33 × median event read count, 3.33 × median event read count]. The values were then normalized by the total event count.

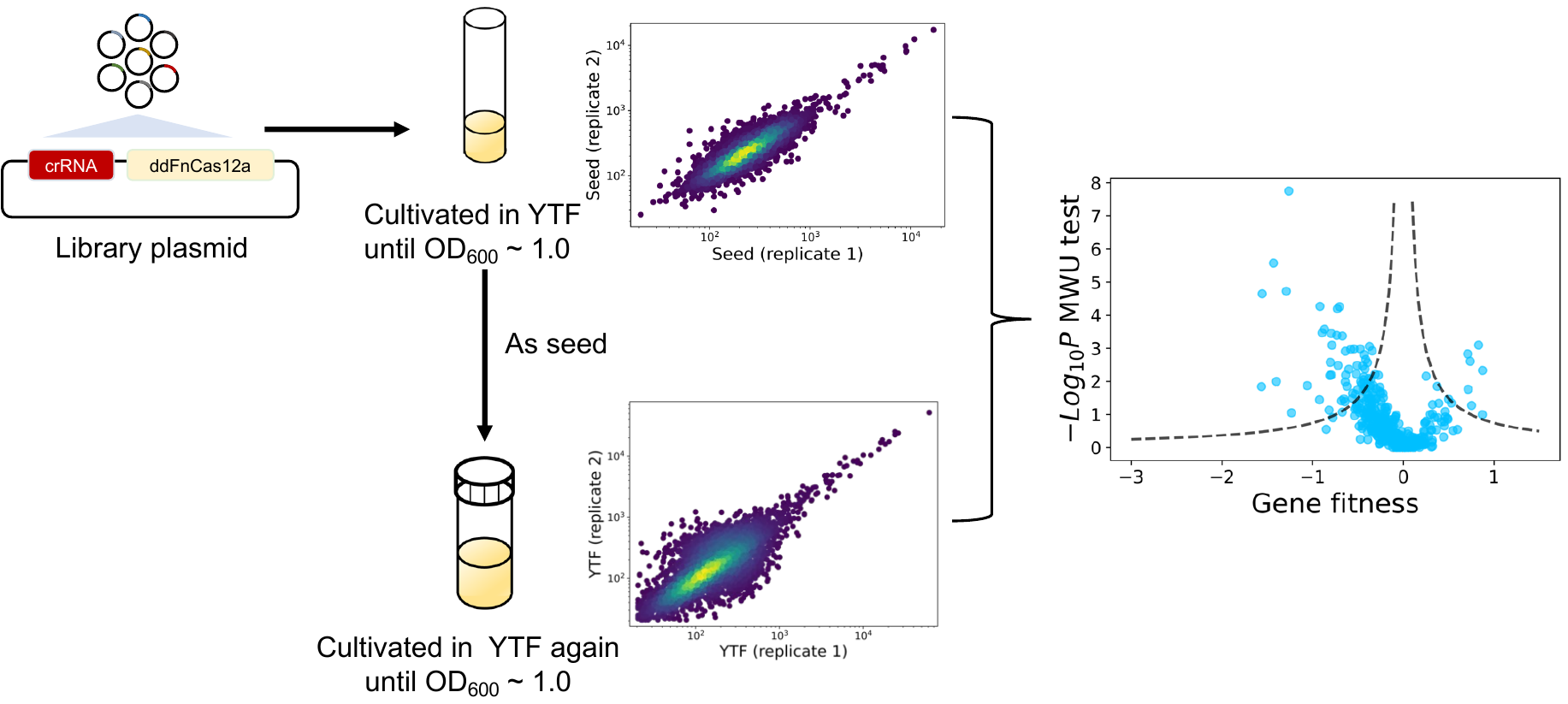

**Supplementary Fig. 2** Flowchart illustrating the screening process for identifying the TF genes associated with the heterotrophic growth of *C. ljungdahlii*.

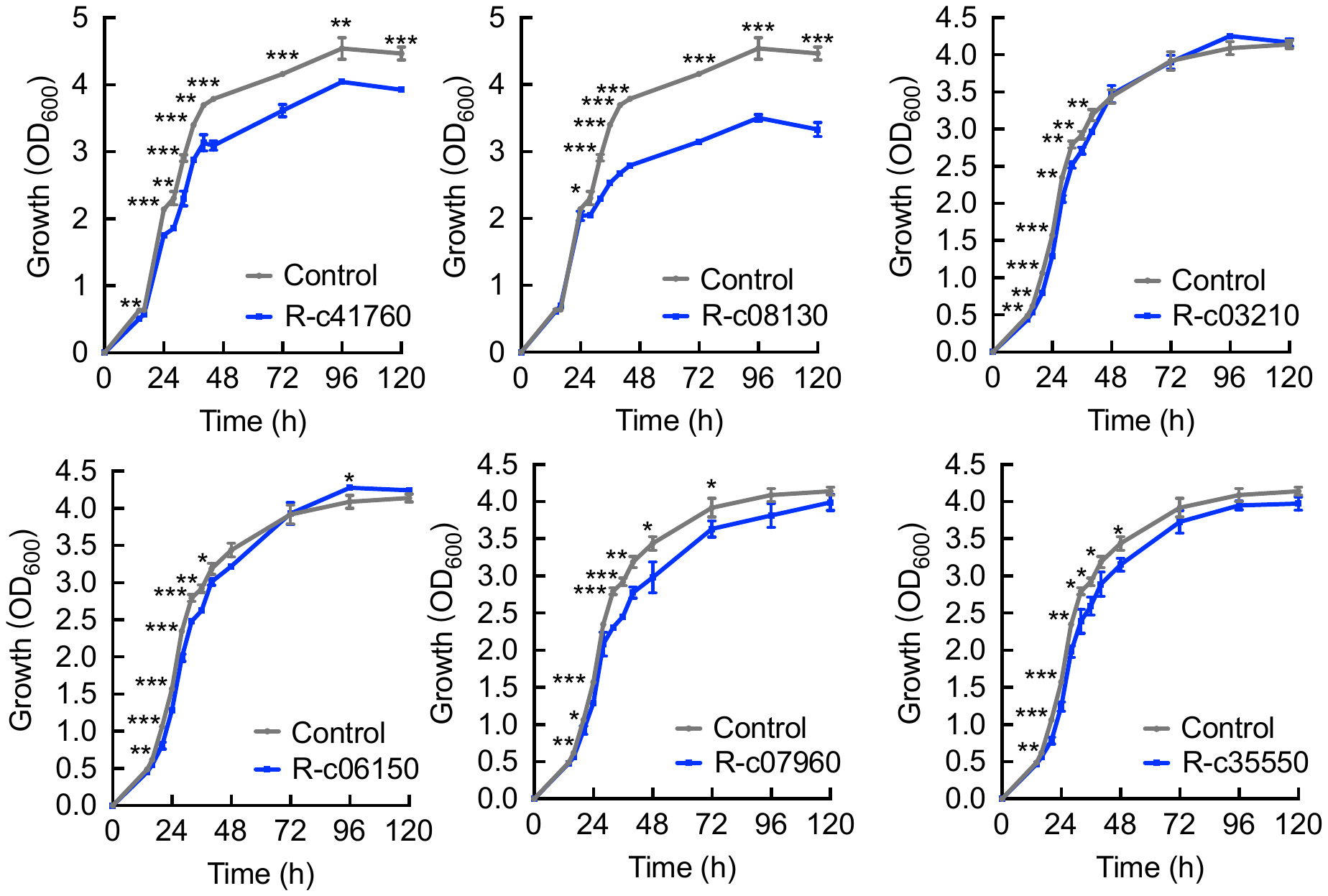

**Supplementary Fig. 3** Influence of the repression of six selected TF genes on the heterotrophic growth of *C. ljungdahlii* in the YTF medium. Data are presented as mean ± standard deviation (*n* = 3). Statistical analysis was performed using the two-tailed Student’s *t*-test. *, *P* < 0.05; **, *P* < 0.01; ***, *P* < 0.001.

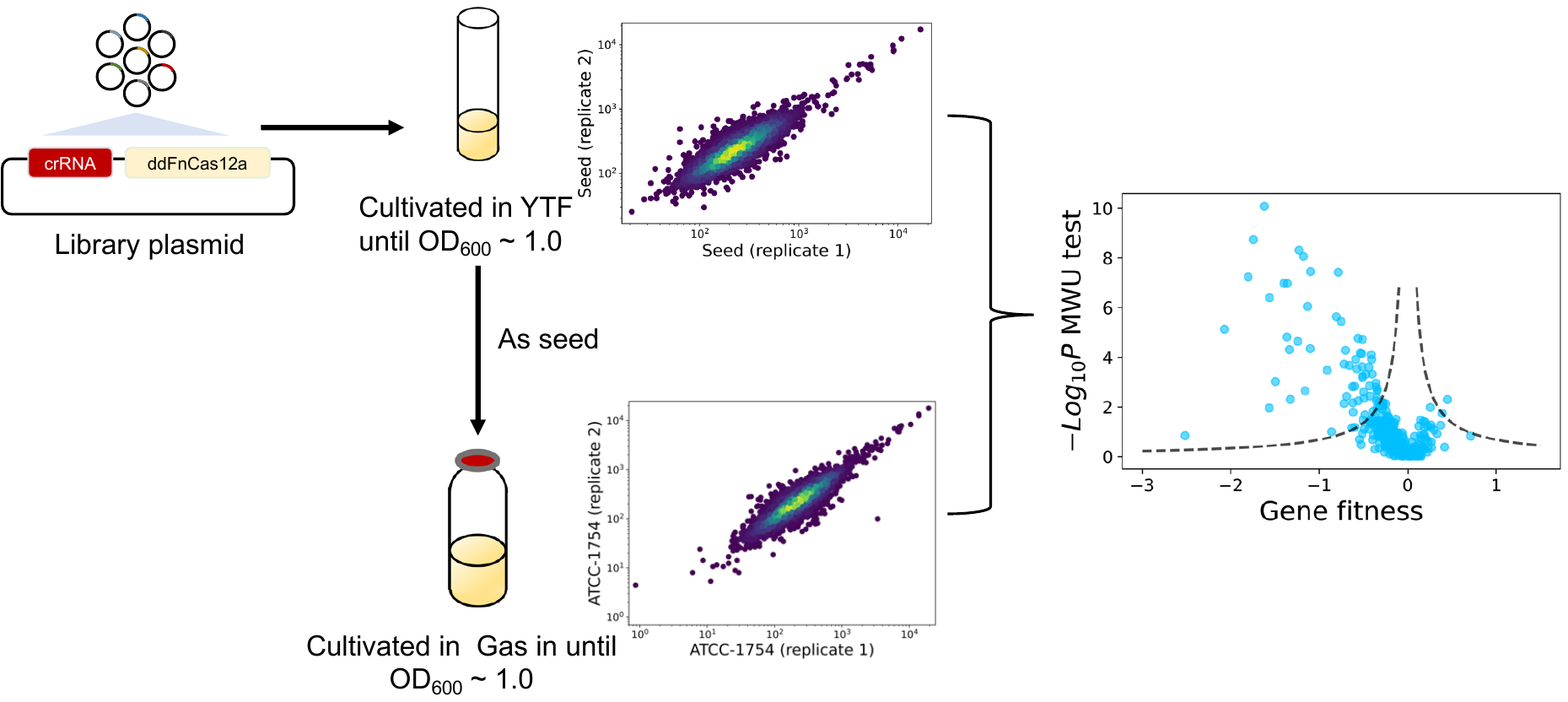

**Supplementary Fig. 4** Flowchart illustrating the screening process for identifying the TF genes associated with autotrophic growth of *C. ljungdahlii i*n gas fermentation.

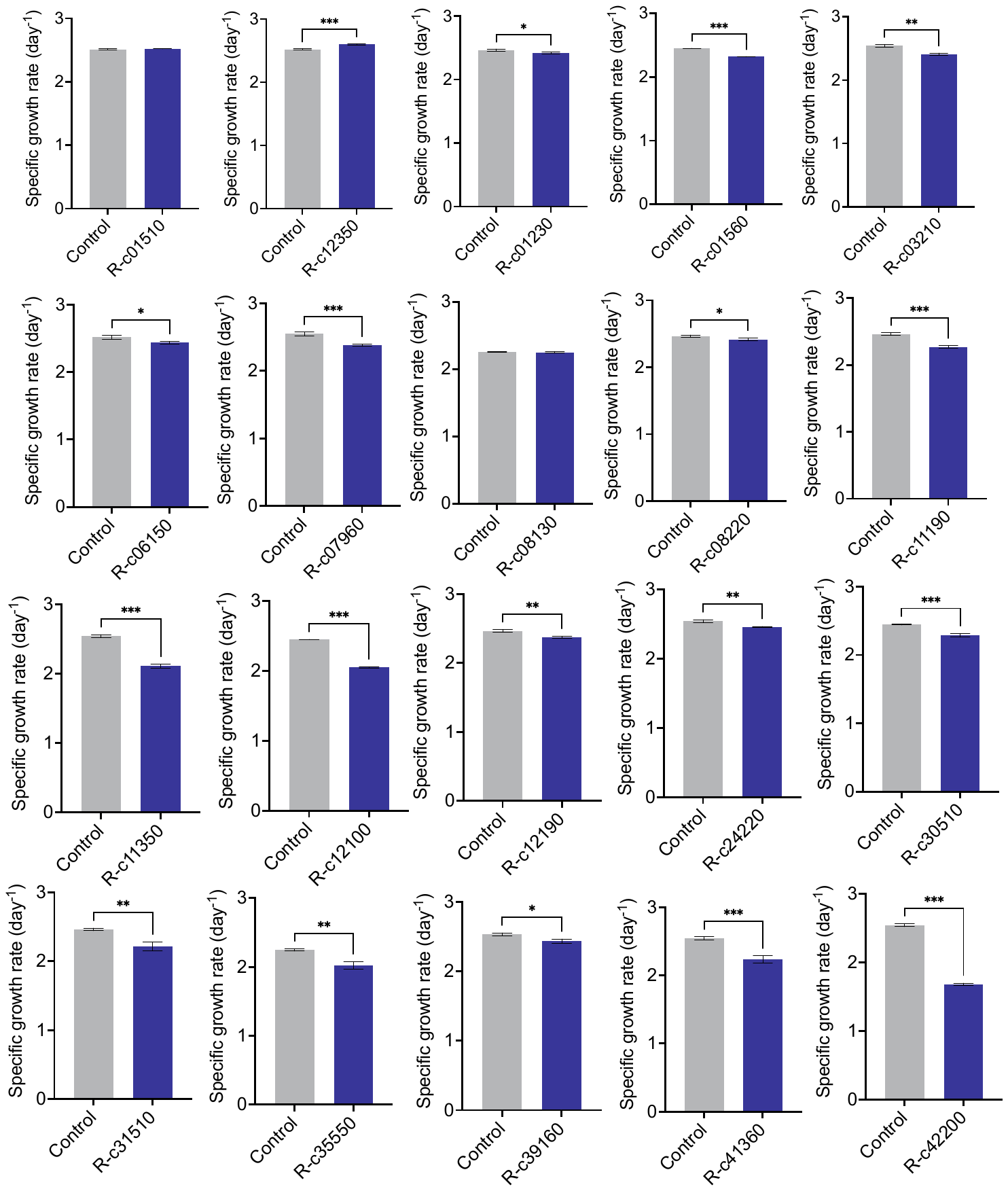

**Supplementary Fig. 5** Influence of the repression of 20 selected TF genes on the growth rate of *C. ljungdahlii* in gas fermentation. Data are presented as mean ± standard deviation (*n* = 3). Statistical analysis was performed using the two-tailed Student’s *t*-test. *, *P* < 0.05; **, *P* < 0.01; ***, *P* < 0.001.

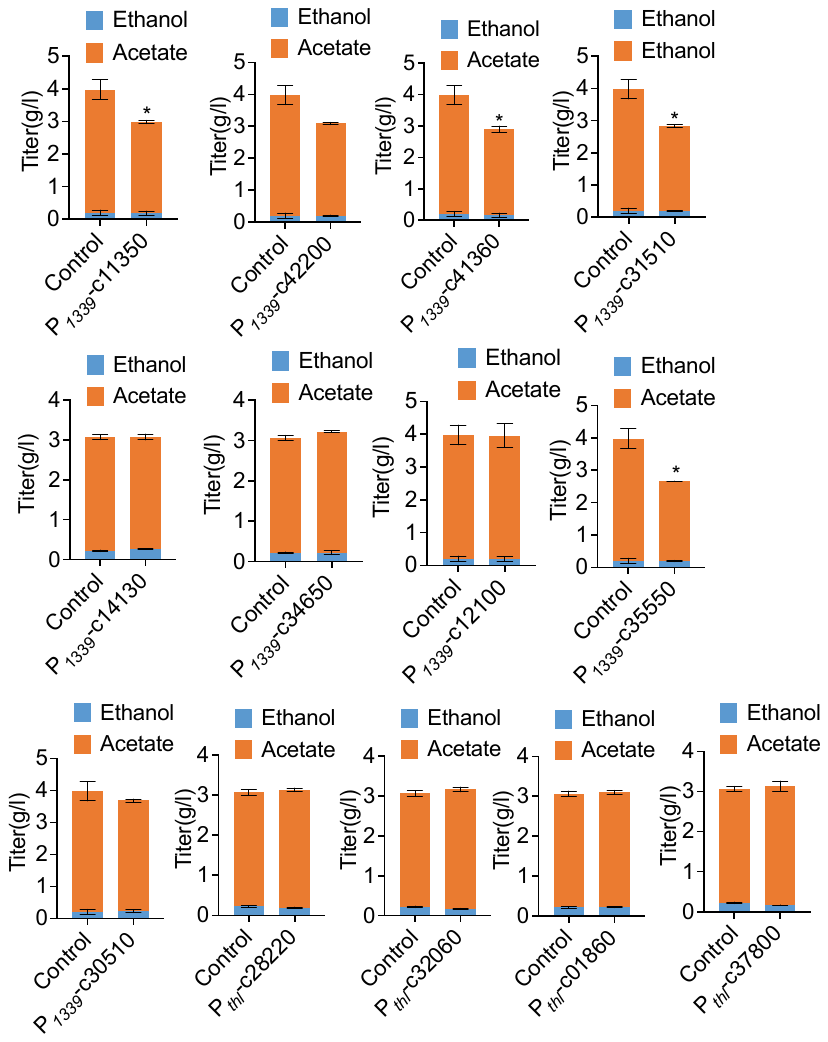

**Supplementary Fig. 6** Influence of the overexpression of 13 selected TF genes on the product (acetate and ethanol) formation of *C. ljungdahlii* in syngas fermentation. Data are presented as mean ± standard deviation (*n* = 3). Statistical analysis was performed using the two-tailed Student’s *t*-test. *, *P* < 0.05.

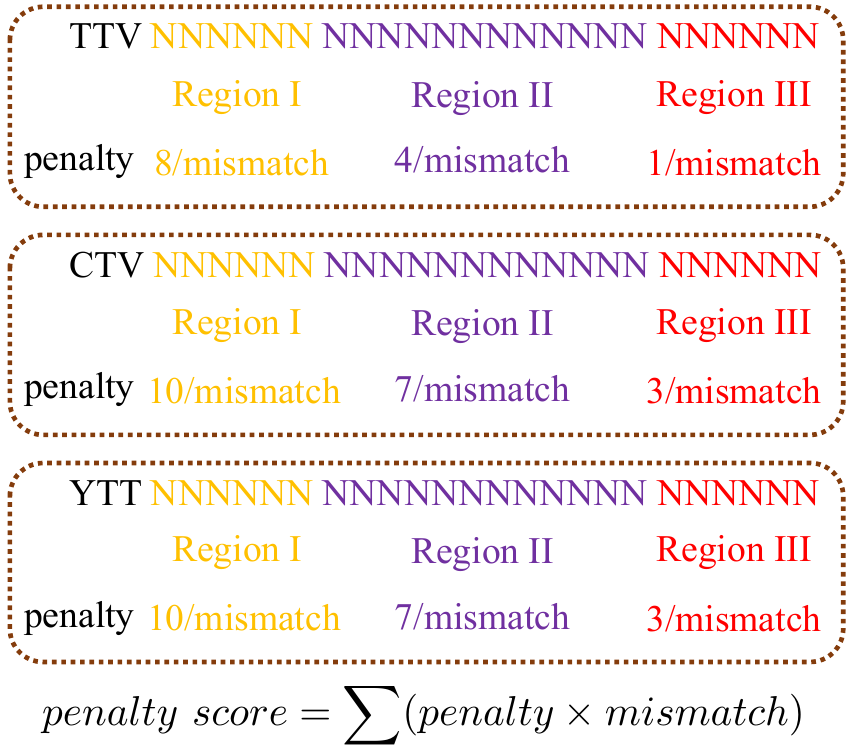

**Supplementary Fig. 7** The scoring matrix used to measure the significance of off-target hits. Potential off-target sites, identified through SeqMap, underwent scoring through the allocation of varying weights to mismatches within three distinct N24 regions. The cumulative weighted sum of all mismatches was then computed and compared to a customized threshold (20). In cases where the sum falls below this threshold, the off-target holds significance, resulting in the removal of the corresponding crRNA.
